# Brewing COFFEE: A sequence-specific coarse-grained energy function for simulations of DNA-protein complexes

**DOI:** 10.1101/2023.06.07.544064

**Authors:** Debayan Chakraborty, Balaka Mondal, D. Thirumalai

## Abstract

DNA-protein interactions are pervasive in a number of biophysical processes ranging from transcription, gene expression, to chromosome folding. To describe the structural and dynamic properties underlying these processes accurately, it is important to create transferable computational models. Toward this end, we introduce **Co**arse grained **f**orce **f**ield for **e**nergy **e**stimation, COFFEE, a robust framework for simulating DNA-protein complexes. To brew COFFEE, we integrated the energy function in the Self-Organized Polymer model with Side Chains for proteins and the Three Interaction Site model for DNA in a modular fashion, without re-calibrating any of the parameters in the original force-fields. A unique feature of COFFEE is that it describes sequence-specific DNA-protein interactions using a statistical potential (SP) derived from a dataset of high-resolution crystal structures. The only parameter in COFFEE is the strength (*λ*_*DNAPRO*_) of the DNA-protein contact potential. For an optimal choice of *λ*_*DNAPRO*_, the crystallographic B-factors for DNA-protein complexes, with varying sizes and topologies, are quantitatively reproduced. Without any further readjustments to the force-field parameters, COFFEE predicts the scattering profiles that are in *quantitative agreement* with SAXS experiments as well as chemical shifts that are consistent with NMR. We also show that COFFEE accurately describes the salt-induced unraveling of nucleosomes. Strikingly, our nucleosome simulations explain the destabilization effect of ARG to LYS mutations, which does not alter the balance of electrostatic interactions, but affects chemical interactions in subtle ways. The range of applications attests to the transferability of COFFEE, and we anticipate that it would be a promising framework for simulating DNA-protein complexes at the molecular length-scale.

**Graphical TOC Entry:** 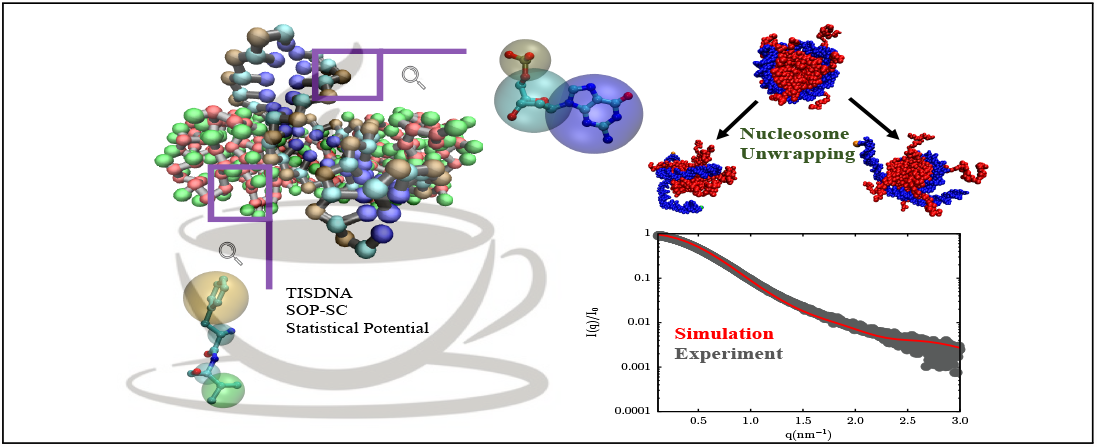

## Introduction

DNA-protein complexes play key roles within the cellular machinery, from orchestrating chromatin organization across different length-scales,^1–3^ to regulating gene expression, replication, as well as DNA repair. Structural studies based on X-ray crystallography, NMR, and most recently cryo-EM techniques show that interactions between DNA and partner proteins could be non-specific, driven largely by electrostatic interactions and size and shape complementarity, or sequence-specific, requiring precise binding modules. Insights into the energetics of DNA-protein interactions have also come from gauging contact statistics within structural databases,^4^ and other knowledge-based approaches. ^5,6^ However, simple rules for predicting the sequence-dependent changes in DNA-protein interactions have not emerged. Indeed, progressing towards a quantitative understanding of the molecular mechanisms governing the assembly of DNA-protein complexes is an important area of research.

Structural studies, although important, provide only a restricted view of the interactions driving the formation of DNA-protein complexes. The dynamics of DNA-protein binding could substantially modulate the efficacy of the recognition process, and tune the desired functionality. For instance, single molecule experiments ^7,8^ as well as computations based on minimal models^9–11^ have revealed that many proteins including RNA polymerases and the tumor suppressor p53, may initially search for target sites on DNA via facilitated diffusion. ^12^

Due to the limited spatial and temporal resolution, experiments alone cannot fully resolve the structural and dynamical details of DNA-protein assembly. Atomically detailed simulations,^13^ based on different force-fields, have provided invaluable insights into various aspects of DNA-protein complexes, including the binding of transcription factors to cognate sites on DNA,^14–16^ the spontaneous association of single-stranded DNA with SSB proteins, ^17^ and salt-dependent unwrapping in nucleosomes.^18,19^ Despite these advances, all-atom simulations of DNA-protein complexes on biologically relevant time-scales are computationally intractable. In addition, force-field deficiencies often manifest themselves only over long simulation time-scales, and may require extensive reparametrization to yield the needed accuracy for characterizing DNA-protein complexes.^20^

It has been shown repeatedly that it is advantageous to exploit a simplified or a ‘coarsegrained’ (CG) representation, especially for large systems such as protein-DNA complexes. By judiciously choosing a resolution that depends on the length and time-scales of interest, insights into a variety of problems could be obtained.^21^ For example, using a single-bead polymer-type model, important predictions have been made regarding the force-induced unwrapping of nucleosomes,,^22^ promoter melting by RNA polymerase,^23^ and more recently the formation of condensates in low complexity RNA sequences. ^24^ In recent years, CG models of different resolutions have been introduced to simulate DNA-protein complexes. Combination of the AWESEM protein force-field^25^ and the 3SPN model for DNA^26^ was used to characterize the energy landscapes for Fis:DNA binding,^27^ as well as nucleosome unwrapping. ^28^ Models of similar resolution have also been exploited by other groups to probe the molecular details of various DNA-protein assembly processes, such as DNA bending induced by architectural proteins,^29^ sliding of ssDNA on protein surfaces, ^30,31^ and higher-order chromatin folding. ^32,33^ In addition, higher resolution CG force-fields such as SIRAH and MARTINI are successful in describing DNA-protein interfaces.^34,35^ Several studies have also exploited a multiscale approach, in which certain regions of the biomolecule, particularly the DNA-protein interfaces are described in atomic detail, and the rest are represented at a CG level. Schulten and coworkers,^36^ and subsequently others, ^37^ have adopted this scheme to probe the conformational dynamics of the lac repressor-DNA complex. However, for models with mixed granularity, the coupling between different resolutions could be non-trivial, which makes it challenging to obtain a balanced description of the energetics.^38^

During the last decade or so, we have introduced several CG models for understanding biomolecular folding and assembly. Among these, the Self-Organized Polymer model with side-chains (abbreviated as SOP-SC) has been remarkably successful in predicting the thermodynamics and kinetics of folding for a diverse range of protein sequences, at different denaturant, salt concentration, as well as pH. ^39–45^ For nucleic acids, the three interaction site (TIS) model^46^ provides a fine balance between accuracy and speed. Different variants of the TIS model have been exploited by us and others^26^ to provide a quantitative description of ion-assisted RNA folding,^47–51^ folding of G-quadruplexes, ^52^ and DNA mechanics and thermodynamics.^53^ Encouraged by the wide range of applicability of our existing CG models, we develop **COFFEE** (**Co**arse-grained **F**orce **F**ield for **E**nergy **E**stimation), a hybrid potential that integrates the SOP-SC model for folded proteins, with the TIS model for DNA. It is worth emphasizing that while making COFFEE, we did not recalibrate the parameters in the SOP-SC or TIS energy functions, which individually provides quantitatively accurate description of the sequence-specific protein-protein and DNA-DNA interactions, respectively. A key ingredient of COFFEE is a knowledge based statistical potential (SP) used to describe the sequence-specific DNA-protein contacts.

We first show that the simulations based on COFFEE accurately reproduce the crystallographic B-factors for a variety of DNA-protein complexes with different sizes and topologies. In addition, COFFEE also predicts the scattering profiles in quantitative agreement with SAXS experiments, and chemical shifts that are consistent with NMR, *without any further fine-tuning*. As a further application of COFFEE, we probe the salt-induced unwrapping of a nucleosome. We show that various partially unwrapped nucleosome conformations are populated as the monovalent salt concentration is increased. Our observations recapitulate the key findings from previous simulations,^54,55^ as well as SAXS experiments. ^56,57^ We also show that ARG to LYS mutations at defined superhelical locations destabilize the nucleosome structure, causing extensive unwrapping, in accord with experimental findings. ^58,59^

The accuracy of our findings attests to the power of COFFEE. We anticipate that the method underlying COFFEE would be a promising framework for simulating large DNA-protein assemblies, particularly at the 10-100 nm length-scale, and would buttress the ongoing efforts^13^ to transform the current state-of-the-art.

## Methodology

COFFEE combines the Self Organized Polymer with Side-Chains (SOP-SC) model for proteins^39,60^ with a Three Interaction Site (TIS) model for DNA.^46,53^ In the SOP-SC model, each amino-acid residue is represented using two interaction sites, a backbone bead (BB) centered on the C_*α*_ atom and a side-chain (SC) bead placed on the center-of-mass of the side-chain. In the DNA model, each nucleotide is represented using three interaction sites, positioned on the center of mass of the phosphate (P), sugar (S), and base (B) groups. The coarse-grained energy function for the DNA-protein complex is:

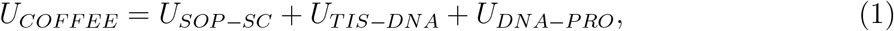

where *U*_*SOP−SC*_ is the energy function corresponding to the SOP-SC model; *U*_*TIS−DNA*_ is the energy function for the TIS-DNA model; and *U*_*DNA−PRO*_ describes the specific as well as non-specific interactions between the DNA and protein molecules.

### SOP-SC model for proteins

The SOP-SC model is optimized for studying single-as well as multi-domain protein folding^61^ under a diverse set of conditions, and describes the thermodynamics and kinetics in quantitative agreement with experiments. ^39,41–45^ The SOP-SC energy function includes contributions from bonded (*U*_*FENE*_), native (*U*_*N*_), non-native (*U*_*NN*_), as well electrostatic interactions 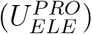, and is given by:

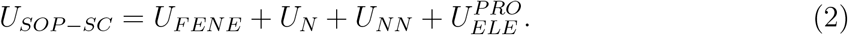

### TIS model for DNA

The TIS model for DNA provides a quantitatively accurate description of both single-stranded and double-stranded DNA, and has recently been used^53^ to explore their sequence-dependent mechanical as well as thermodynamic properties. The TIS energy function includes contributions from bond (*U*_*B*_), angular (*U*_*A*_), stacking (*U*_*S*_), hydrogen-bonding (*U*_*HB*_), excluded-volume (*U*_*EV*_), and electrostatic 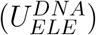 interactions:

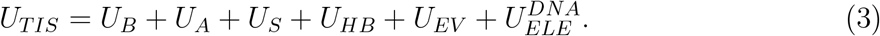

The detailed functional forms of the potentials in Eq. 2, and 3 are described in the Supporting Information (SI). The force-field parameters for the SOP-SC and TIS models are tabulated in the SI (Tables S1 to S7).

### DNA-protein interactions

The non-bonded interactions between DNA and proteins, as described by the *U*_*DNA−PRO*_, includes both electrostatic and non-electrostatic components,

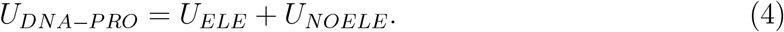

The electrostatic interactions, *U*_*ELE*_, between the phosphates on DNA and the amino-acid side-chains of the protein are described by the Debye-Hückel potential,

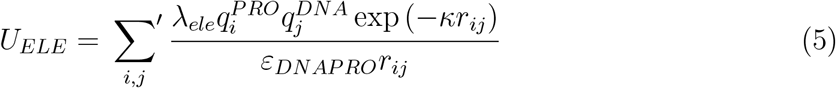

where 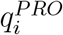 is the charge on the SC bead of amino-acid residue *i* (+1 for ARG and LYS, and -1 for ASP and GLU at physiological pH), and 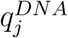 denotes the renormalized charge (chosen to be 0.6 to account for counter-ion condensation, ^62^ as described previously.^53^) on the phosphate bead corresponding to nucleotide *j*; *r*_*ij*_ is the distance between the SC and the phosphate beads; *κ* denotes the inverse Debye length, and *ε*_*DNAPRO*_ is the dielectric constant. We assume that *ε*_*DNAPRO*_ to be temperature-independent, and set it equal to 78.0. Following previous studies,^28,63,64^ we also include a scaling factor, *λ*_*ele*_=1.67, to partially account for the lack of explicit counter-ions in the model.

The non-electrostatic component, *U*_*NOELE*_, includes both native (*U*_*N*_) and non-native (*U*_*NN*_) contributions,

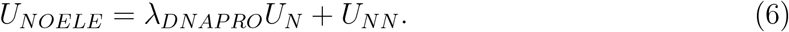

In Eq. 6, the adjustable parameter *λ*_*DNAPRO*_ tunes the strength of the native interactions in the DNA-protein complex. A contact is presumed to be native if the distance between a DNA and a protein bead in the reference structure is less than 12 Å. The native interactions among the DNA and protein beads are described in terms of Lennard-Jones type potentials:

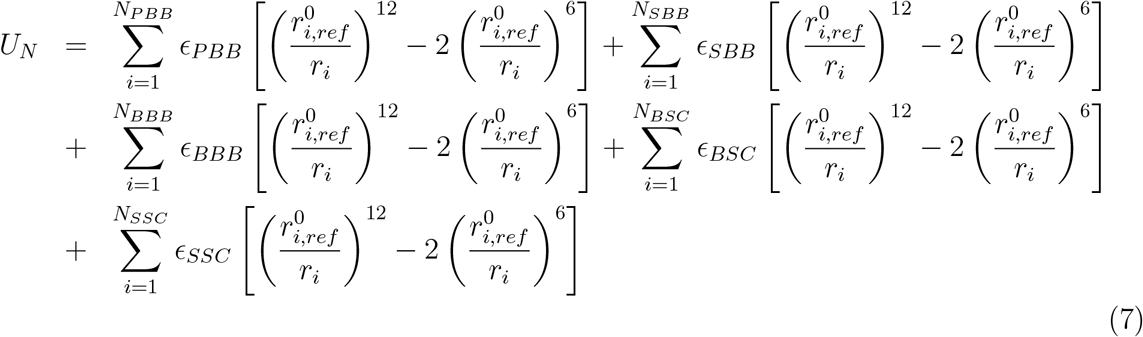

The first term in Eq. 7 accounts for the native interactions between the phosphates (P) of DNA and the protein backbone (BB), with *ϵ*_*PBB*_ being the interaction energy scale. The second and the third terms denote the pairwise interactions between the protein backbone (BB) with the sugars (S) and nucleobases (B) of DNA. The strengths of these non-bonded interactions are denoted by *ϵ*_*SBB*_ and *ϵ*_*BBB*_, respectively. Based on the distance cutoff, native contacts can also be defined between the protein side-chains (SC) and the DNA nucleobases (B), as well as sugars (S). The corresponding interaction energies are denoted by *ϵ*_*SSC*_ and *ϵ*_*BSC*_. Following previous works,^65,66^ we assume that the interactions between the DNA phosphate backbone and the amino-acid side-chains is purely electrostatic in nature. In all cases, 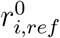 denotes the distance between the *i*^*th*^ pair of DNA and protein beads in the coarse-grained representation of the reference (experimental) structure.

The energy scales for the *k*^*th*^ pair of native interactions, depend on the protein and DNA sequences, and can be generically represented as, 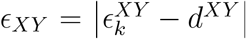, where *X* ∈ [*P, B, S*] denotes the type of DNA bead, and *Y* ∈ [*BB, SC*] denotes the type of protein-bead. Here, *ϵ*^*XY*^ is related to the ‘free energy’ cost, Δ*G*^*XY*^, of forming a contact between residues *X* and *Y*, and *d*^*XY*^ is an offset parameter, which ensures that *ϵ*_*XY*_ values are positive. The 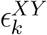 values for the different amino-acid-nucleotide pairs were derived from the statistics of DNA-protein contacts, as described below.

### Statistical potential for native DNA-protein contacts

The development of statistical potentials (SPs) based on residue-residue contacts found within structures deposited in the PDB database, was pioneered by Tanaka and Scheraga^67,68^ in their studies on protein folding. Subsequently, various improvements were suggested by Miyazawa and Jernigan, ^69^ and others.^70–72^ The concept of SPs was also extended to RNA folding, and used for RNA secondary structure prediction. ^73^ To generate the contact statistics, we consider the non-redundant set of DNA-protein complexes available in the Protein-DNA Interface Database.^74^ The PDB IDs of the complexes are listed in the Supporting Information (Table S8). The amino-acid-nucleotide contact maps were computed after coarse-graining the atomistic structure of each DNA-protein complex.

The generation of SPs for DNA-protein contacts involves the following steps. (i) Let 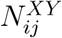 be the total number of amino-acids of type *i* in contact with nucleotide of type *j* within the non-redundant set of PDB structures. The superscript *X* ∈ [*P, B, S*] denotes type of the DNA bead, and *Y* ∈ [*BB, SC*] denotes the type of protein bead. For a contact to exist between *X* and *Y*, the distance of separation must be less than or equal to 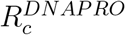. (ii) The probability, 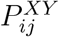, that a nucleotide *i* is in contact with an amino-acid *j* is given by: 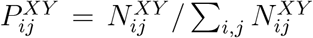. (iii) In the computation of SPs, the choice of the reference state is important.^73^ For simplicity, we assume the random occurrence of a pair *ij* to be the reference state. The probability that a pair *ij* would be in contact by random chance within the ensemble of PDB sequences is given by: 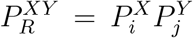. (iv) Assuming that the ensemble of structures deposited within the PDB database are at quasi-equilibrium, and that Boltzmann statistics applies, the effective contact ‘free energy’ (or equivalently the SP), 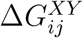, between nucleotide *i* and amino-acid *j* is given by:

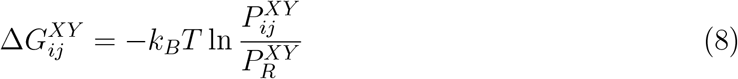

where *k*_*B*_ is Boltzmann’s constant, and *T* is the effective temperature. The interaction energy 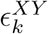 for the *k*^*th*^ pair (between nucleotide *i* and amino-acid *j*) is simply 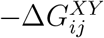. Despite well-known limitations of this approach,^75,76^ SPs have been used successfully in applications related to biomolecular structure prediction,^67,73^ thermodynamics and dynamics of protein folding, ^41,43^ and more recently in simulations of chromosome organization.^77^

The 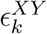 values estimated from the contact statistics are tabulated in the Supporting Information (Table S9 to S13). As an illustrative example, we show the variations in the 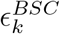 values for contacts between different nucleobases (B) and amino-acid side-chains (SC) in Figure 1. We find that Guanine-Arginine contacts are the most favorable, whereas Guanine-Leucine contacts are least favored, in agreement with a previous statistical survey of DNA-protein interactions.^78^

**Figure 1:**
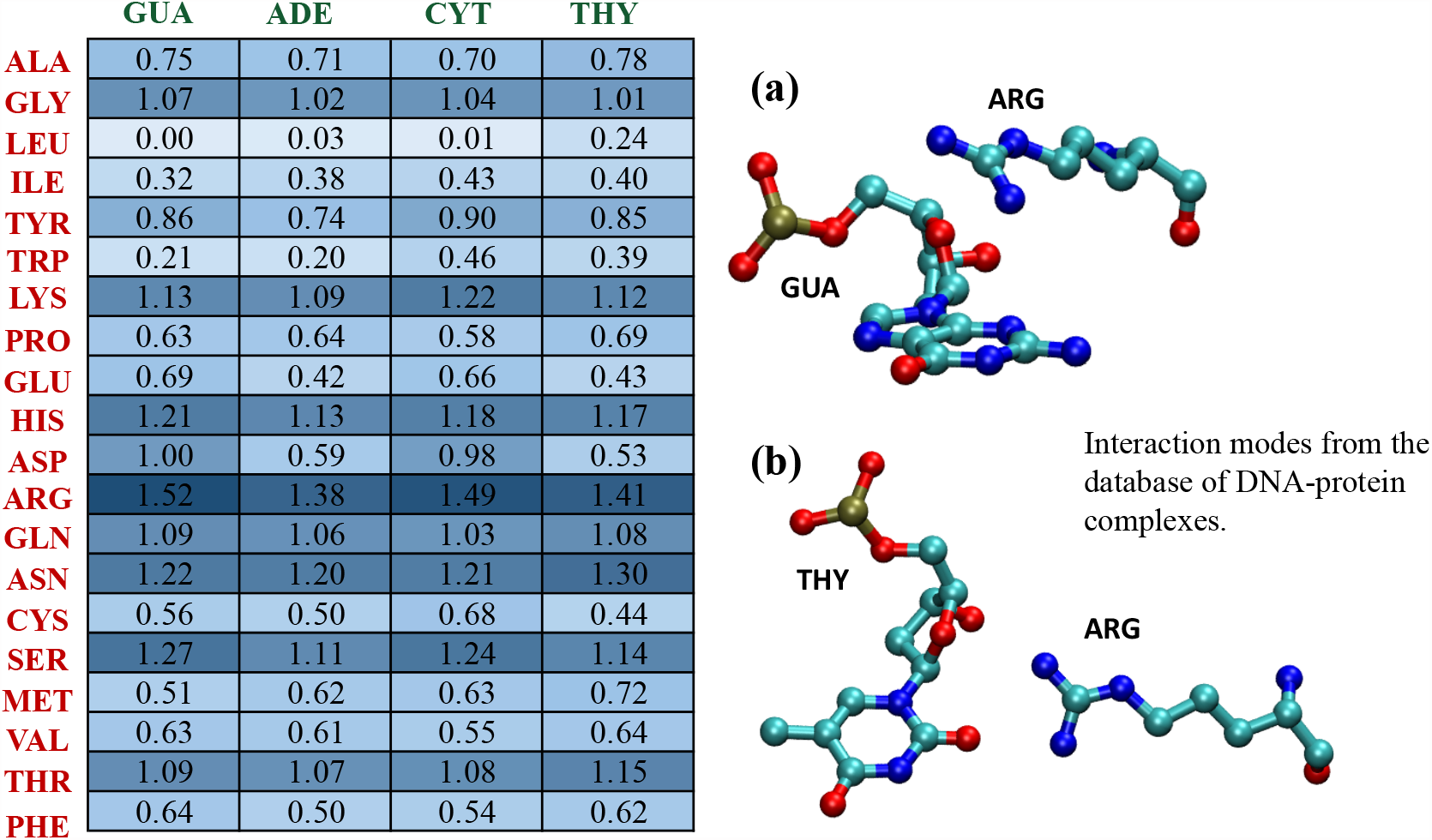
Left: A matrix denoting the effective energy scales (in kcal/mol) derived from the statistics of DNA-protein contacts for nucleobases (B) and amino-acid side-chains (SC). Each cell of the matrix is color-coded in accordance with the strength of the DNA-protein contact, with intense colors denoting stronger interactions. Right: Illustrative examples of contacts involving (a) GUA and ARG; (b) THY and ARG.

The non-native interactions, 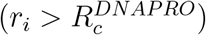 between DNA and protein are taken to be purely repulsive. The interaction potential, *U*_*NN*_ is given by,

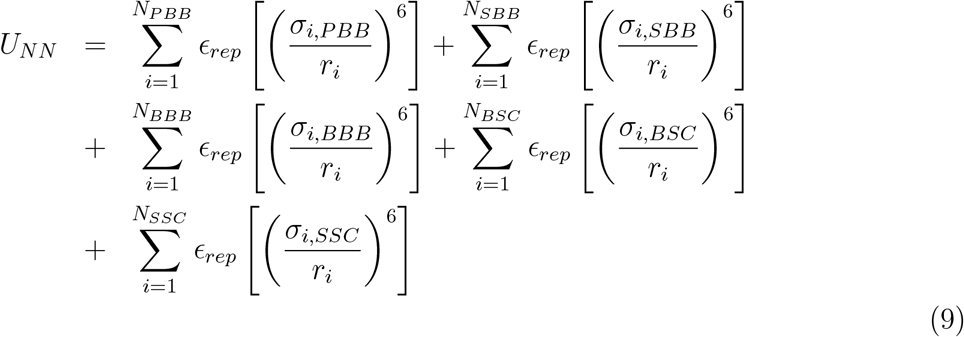

In the above equation, 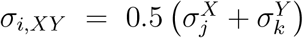 is the van der Waals radius for the *i*^*th*^ nucleotide-amino-acid pair; 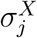 and 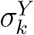 denote the radii of DNA bead *j* of type *X*, and protein bead *k* of type Y, respectively. The value of *ϵ*_*rep*_ is set to 1 kcal/mol. The van der Waals radii for the different DNA and protein beads are tabulated in the Supporting Information (Tables S2 and S5).

### Calibrating the DNA-protein contact potential

The only free parameter in the COFFEE is *λ*_*DNAPRO*_ (Eq. 6), which tunes the overall strength of the native interactions. To calibrate the contact potential, we initiated simulations from a high-resolution crystal structure of the human TBP core domain complexed with DNA (PDB ID: 1CDW), ^79^ for different values of *λ*_*DNAPRO*_. Beyond *λ*_*DNAPRO*_ *≈* 0.15, there is no significant change in the root-mean-squared (RMSE) between the B factors reported in the crystal structure, and those calculated from our simulations (Fig. 2(a) and 2(c)). For *λ*_*DNAPRO*_ = 0.15, the residue-specific B factors estimated from simulations are also in good agreement with experimental values (Fig. 2(b) and 2(d)). However, for some residues, the fluctuations are enhanced, which is expected because our simulations do not strictly model the crystal environment.

**Figure 2:**
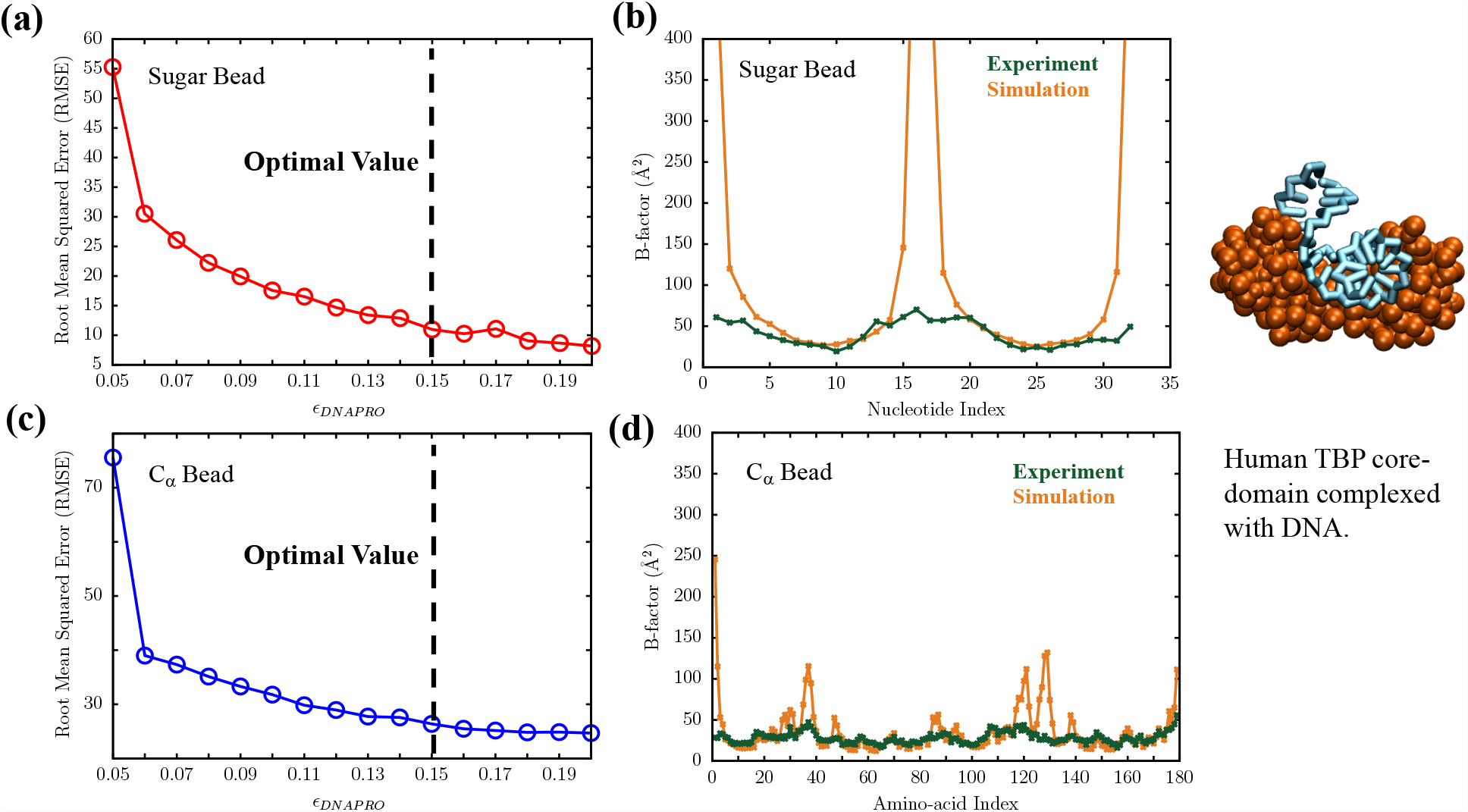
Calibration of the DNA-protein contact potential using a high-resolution crystal structure of the human TBP-core domain in complex with DNA. The DNA duplex is shown in cyan, and the TBP domain is rendered in brown using a space-filling representation. (a) The variation of the Root Mean Squared Error (RMSE) between experimental and simulated B-factors for the sugar bead. (b) Comparison between the crystallographic B-factors (green) and those estimated from COFFEE simulations (orange) for the sugar groups.(c) The variation of the RMSE between experimental and simulated B-factors for the C_*α*_ bead. In (a) and (c), the dashed line indicates the optimal value of *ϵ*_*DNAPRO*_ = 0.15. (d) Comparison between the crystallographic C_*α*_ B-factors (green) and those estimated from simulations (orange).

### Simulations

To efficiently sample the conformational space of DNA-protein complexes, we carried out Langevin Dynamics (LD) simulations in the underdamped regime. The equation of motion for bead *i* is, 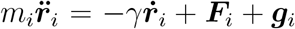, where *m*_*i*_ denotes the mass of the bead, *γ* denotes the frictional drag coefficient, ***F***_*i*_ denotes the conservative force that acts on bead *i* as a result of interactions with other beads; ***g***_*i*_ is a Gaussian random force, which satisfies ⟨ ***f***_*i*_(*t*)***f***_*j*_(*t*^*′*^)⟩ = 6*k*_*B*_*Tγδ*_*ij*_*δ*(*t − t*). Each simulation was carried out at 298 K for 10^8^ steps. Simulations were initiated from twenty different random number seeds to obtain meaningful statistics for any observable. In addition to a CPU implementation within our in-house code, we also ported COFFEE to OpenMM,^80^ and exploited its optimal performance on GPUs to accelerate some of the nucleosome simulations.

### SAXS profiles

The SAXS profiles for the DNA-protein complexes were computed from the trajectories using the Debye formula,

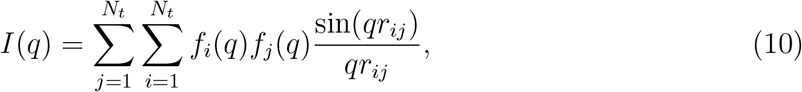

where *f*_*i*_(*q*) and *f*_*j*_(*q*) are the *q*-dependent coarse-grained form factors for beads *i* and *j*, respectively. The residue-specific form factors for the protein backbone and different amino-acid side-chains were taken from Table S2 of Tong *et al*.^81^ The nucleotide-specific coarse-grained form factors were derived using the independent bead approximation ^82^ from a database of high-resolution DNA crystal structures. Further details of this procedure are included in the Supporting Information.

### Calculation of Chemical Shifts

The C_*α*_ chemical shifts were calculated from the trajectory using the LARMOR-C_*α*_ formalism. ^83^ This method exploits a number of geometrical features based on C_*α*_-C_*α*_ distances, and is trained using a random forest classifier on the RefDB database. As shown previously,^83^ LARMOR-C_*α*_ predicts chemical shifts in quantitative agreement with experiment, for a number of folded proteins.

### Calculation of the number of unwrapped base-pairs

To probe the extent of nucleosome unwrapping at different salt concentrations, we use an order parameter introduced previously.^27^ For each base pair *b*, we determined if it is bound to the histone core using:

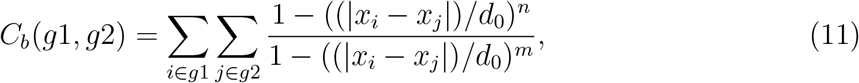

where the switching function 1 *−* ((|*x*_*i*_ *− x*_*j*_|)*/d*_0_)^*n*^*/*1 *−* ((|*x*_*i*_ *− x*_*j*_|)*/d*_0_)^*m*^ is bound between 0 and 1; group *g*1 includes the sugar beads of the base-pairs, and *g*2 includes the *C*_*α*_ atoms of the histone core proteins. Following a previous study,^27^ we set *d*_0_ = 10 Å, *n* = 6 and *m* = 12. *C*_*b*_ is rescaled to lie between 0 and 1 using another switching function, ^27^

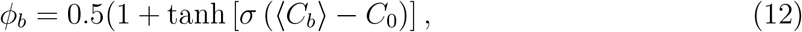

where the angular brackets denote thermal averaging, *σ* = 4.0 and *C*_0_ = 1.5.

The total number of unwrapped base pairs is given by:

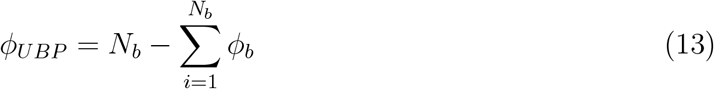

where *N*_*b*_ is the total number of base-pairs.

## Results and Discussion

### COFFEE reproduces the crystallographic B-factors for diverse DNA-protein complexes

To assess the transferability of COFFEE, we simulated six DNA-protein complexes with different sizes, topologies, and number of subunits. In all cases, the simulations were initiated from the crystal-structures deposited in the PDB database (see below).

### Yeast TBP/TATA-box complex

In the crystal structure (PDB ID: 1YTB), a sharp bend near the major groove of the hairpin facilitates the binding between the TATA-box and the TATA-box binding protein (TBP).^84^ The DNA-protein interface in this complex is primarily stabilized by hydrophobic contacts, which remain stable during the course of our simulations. As shown in Fig. 2(a), crystallographic B-factors for the sugar moieties, and the C_*α*_ beads of amino-acid residue are accurately reproduced by COFFEE. In the simulations, the DNA ends are unconstrained, and often fray along the trajectory, resulting in relatively high B-factors for the terminal residues. Some of the loop residues connecting different *β*-strands of TBP exhibit enhanced fluctuations, and are associated with higher B-factors (residues *≈* 34-40 and 126-130). This is likely due to the absence of crystal packing forces, which stabilize the crystal structure, resulting in suppressed motion.

### TFIIB/TBP/TATA-element ternary complex

The high-resolution crystal structure (PDB ID: 1VOL) of the complex formed between human transcription factor IIB (TFIIB), TBP and the TATA element provides structural insights into the early steps of transcription initiation.^85^ In TFIIB, a two-domain *α*-helical protein establishes contacts with the TBP and the TATA element through extensive protein-protein and DNA-protein interactions. The B-factors of the TATA element are well reproduced by our simulations (Fig. 2(b)). The ordered secondary structure elements (helical motifs and *β*-strands) of the TFIIB and TBP domain are stable throughout the trajectory, with the C_*α*_ fluctuations that are comparable to the crystallographic B-factors. On the other hand, disordered loops connecting the different ordered segments, as well some of the terminal residues exhibit enhanced fluctuations. This is reflected in the relatively high B-factors in multiple stretches of residues (for example, *≈* 189-212 and 238-250).

### Brf2-TBP complex bound to DNA

The Brf2-TBP complex bound to its natural U6 promoter (PDB ID: 4ROC) provides a molecular view of transcriptionally active pre-initiation complexes. ^86^ The DNA-TBP binding interface is similar to that in the TFIIB/TBP/TATA-element complex, and many studies have linked this striking resemblance to a common architectural design of the core in different initiation complexes of the transcription machinery.^87^ Just as in the previous two examples, the B-factors of the DNA strands are quantitatively reproduced. In this ternary complex, the different *α*-helices within the cyclin repeats of Brf2 are connected by flexible loops. Several residues within these loops (*≈* 115-132 and *≈* 213-226) are associated with high B-factors. The disordered linker connecting the TBP anchor domain and a Brf2-specific small helical motif (termed as the “molecular pin”) remains highly dynamic during the simulations, with a long stretch of residues (*≈* 250 to 303) exhibiting significant deviations from the crystal structure.

### Engineered I-CreI derivative in complex with DNA from XPC gene

In this engineered complex (PDB ID: 2VBJ), the DNA-protein interface is formed between a heterodimeric derivative of I-CreI (termed as Amel3-Amel4) and 24 bp DNA from the XPC gene.^88^ I-CreI is a member of the homing endonuclease family, and because of its high specificity, it can selectively cleave DNA sequences in complex genomes. ^88,89^ The B-factors of the sugar moieties computed from our simulations are somewhat smaller than those reported for the crystal structure, and do not exhibit position-dependent variations (Fig. 3(d)). This could imply that our coarse-grained simulations do not fully capture the nuanced features of DNA recognition that make I-CreI and its derivatives highly specific scaffolds for genome manipulation. The Amel3-Amel4 heterodimer consists of different secondary structure motifs (*α*-helices and *β*-strands), which are connected by short loops. As is evident from Fig. 2(d), the crystallographic B-factors of the ordered elements are quantitatively reproduced in our simulations. Unlike the other examples that we have considered, in this complex close-range contacts at the DNA-protein interface also involve many residues within the loops. Hence, it is not surprising that loop residues only exhibit minimal fluctuations along the trajectory (Fig 2(d)), and stay close to their crystallographic coordinates.

**Figure 3:**
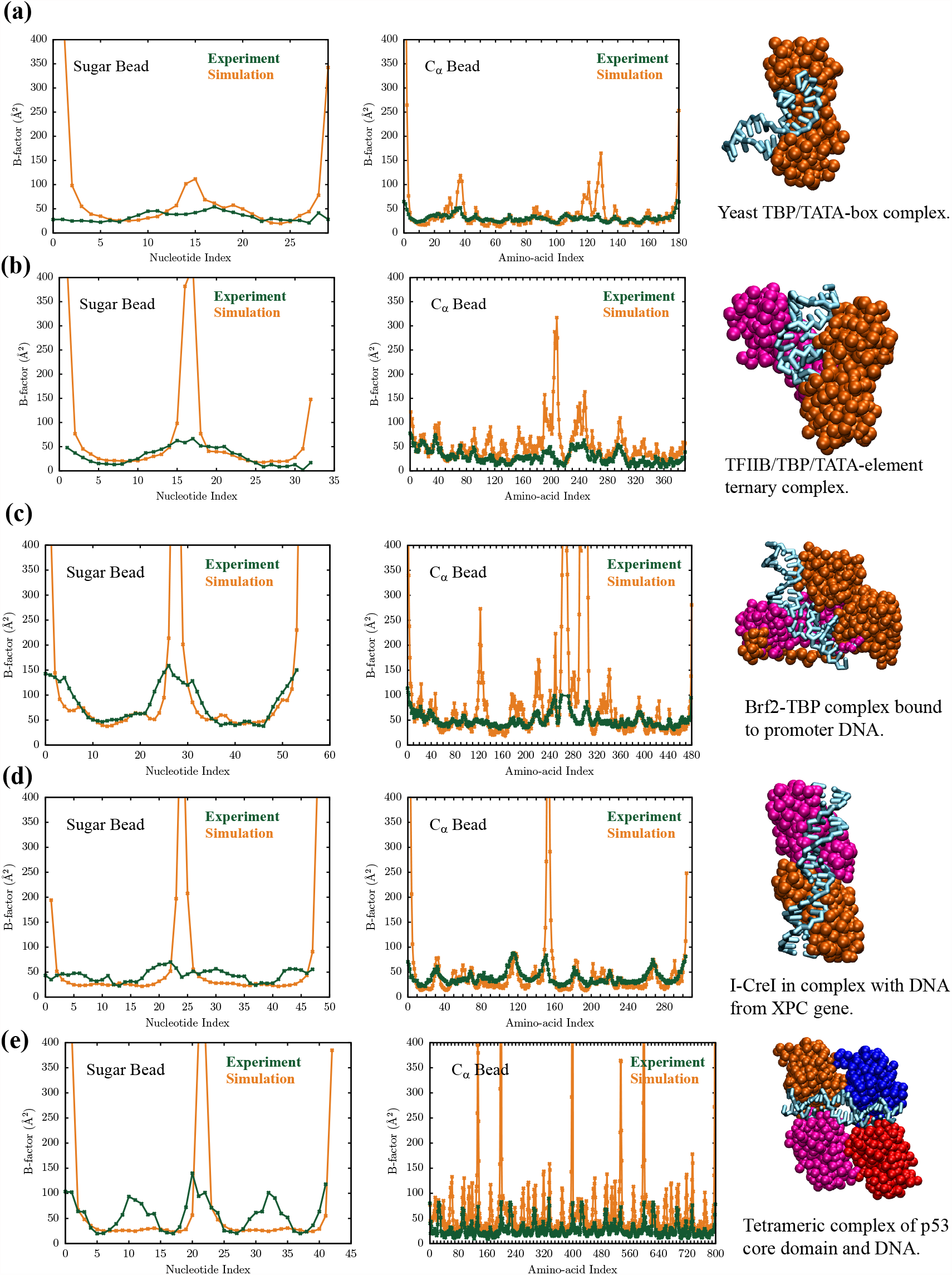
Comparison of the crystallographic B-factors (green) with estimates from COFFEE simulations (orange) for (a) Yeast TBP/TATA-box complex (PDB ID: 1YTB). (b) TFIIB/TBP/TATA element ternary complex (PDB ID: 1VOL). (c) Brf2-TBP complex bound to DNA (PDB ID: 4ROC). (d) I-CreI in complex with DNA from the XPC gene (PDB ID: 2VBJ). (e) Tetrameric complex of p53 with DNA (PDB ID: 4HJE). A coarse-grained representation of each complex is shown on the right. The DNA molecule is shown in cyan, and the individual protein subunits are rendered in different colors using a surface-filling representation.

### Tetrameric complex of p53 with DNA

The crystal structure of the tetrameric core domain of p53 bound to a 21 bp response element (RE) from the BAX promoter gene (PDB ID: 4HJE) is somewhat unusual because unlike other p53-RE complexes, ^90^ a single bp spacer (G11-C32) is embedded within the middle of the DNA sequence. ^91^ The spacer induces local distortions within the DNA duplex, resulting in enhanced B-factors for the adjacent basepairs (Fig 3(e), green profile). Our simulations, however, do not recapitulate this trend (Fig 3(e), orange profile), and the B-factors of all nucleotides are practically similar. Chen and coworkers ^91^ argued that the enhanced fluctuations of nucleotides near the spacer could reflect deviations from the canonical Watson-Crick geometry, or transient base-flipping. These transitions are rather slow, occurring on the millisecond time-scale,.^92,93^ Our simulations primarily probe fluctuations around the native basin, and are unlikely to capture such rare events. The residue-wise B-factor of the p53 core are predicted to be consistently higher than experiment (Fig 2(e)). Similar to previous examples, loop residues connecting different secondary structure elements (for example, residues *≈* 197-201), such as *β*-strands and *α*-helices, as well as terminal residues (*≈* 397-400) exhibit the highest fluctuations.

Previous works^94,95^ suggest that a rigorous comparison to experimental B-factors entails a precise description of the crystal lattice as well as the buffer. Although these crystallization conditions were not explicitly taken into our simulation setup, the B-factor estimates are in near quantitative agreement with those reported in the crystal structures, for most, but not all, of the examples. This is encouraging, and attests to the efficacy of our brewing protocol, which combines two independently developed CG models for proteins and DNA, with a DNA-protein interaction potential derived from contact-statistics.

### Calculated scattering profiles are in quantitative agreement with SAXS experiments

Different variants of small-angle X-ray scattering (SAXS) are routinely used to probe the global dimensions of DNA-protein complexes, as well as characterize their structures at low-resolution. ^96,97^ We simulated three DNA-protein complexes of different sizes and topologies: the immunity repressor-DNA complex (209 residues) in which the DNA-protein binding is asymmetric and is mediated by two independent domains,^98^ the PaHigA-DNA complex (904 residues) in which helix-turn-helix motifs from the protein dimers insert into the DNA major groove to form the binding interface,^99^ and the BusR-promoter (910 residues) complex where the binding to a 22 bp DNA duplex is mediated by a coiled-coil tetramer.^100^ As shown in Fig. 4, the computed scattering profiles, *I*(*q*), as a function of the scattering vector *q* are in quantitative agreement with those reported by SAXS experiments. The scattering curves in the Kratky representation (*q*^2^*I*(*q*)*/I*(0) vs *q*) are included in the Supporting Information. In particular, the agreement is remarkable in the low *q* regime (*q* ⩽ 1.3 nm^*−*1^), suggesting that COFFEE accurately reproduces the global dimensions of the DNA-protein complexes. The deviations from the experimental SAXS curves are rather modest even at high *q* values, which shows that our model accurately captures the structural details even at small length-scales. The radii of gyration calculated from simulations 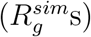 are practically indistinguishable from the values 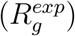 reported from a Guinier analysis of the experimental SAXS profiles (Fig 4). This is striking because we did not tweak any parameter in COFFEE to obtain the reported accuracy of *I*(*q*) in Fig 4.

**Figure 4:**
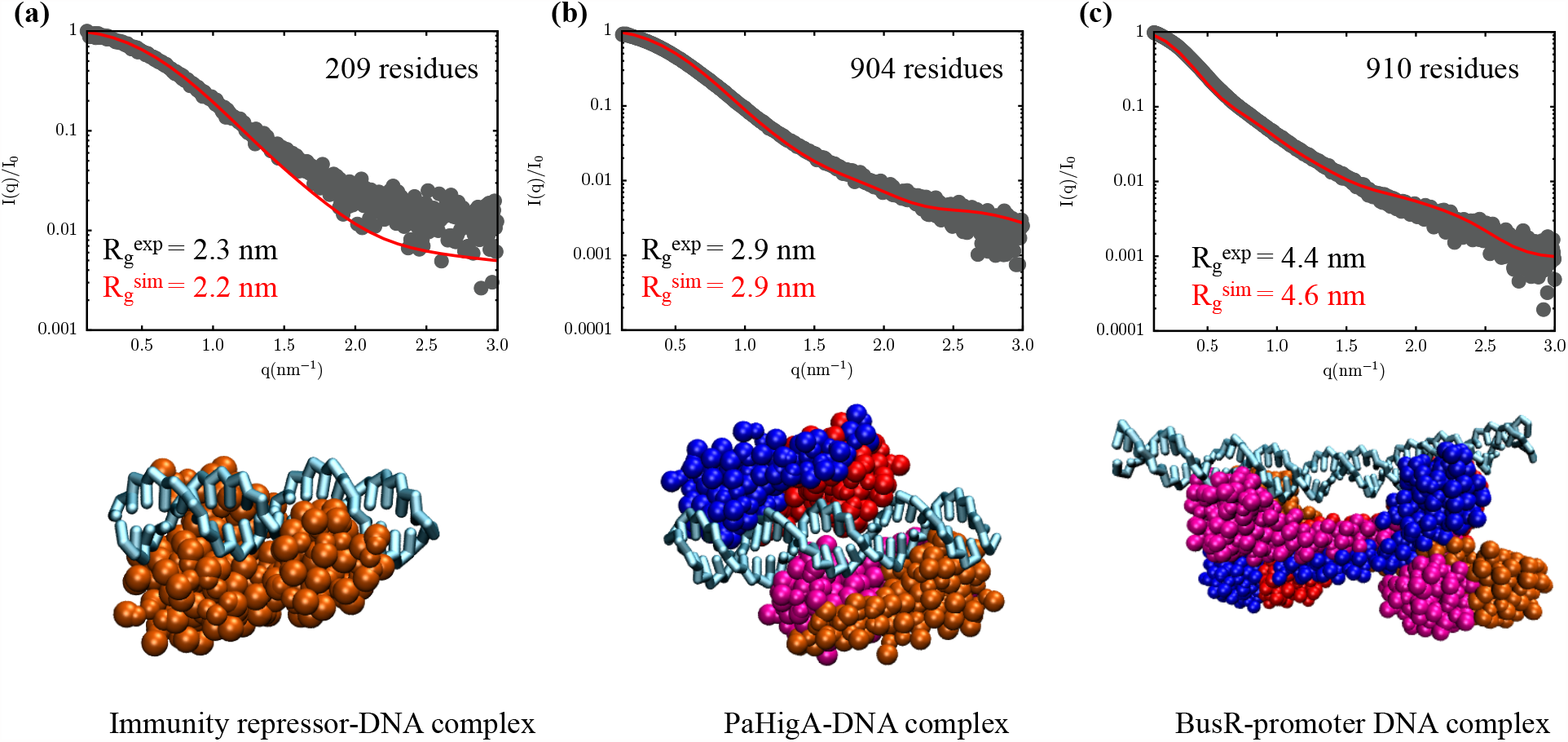
Structure factor calculated from simulations (red curve) and from SAXS experiments (grey points) for (a) Immunity repressor-DNA complex (PDB ID: 7R6R). (b) PaHigA-DNA complex (PDB ID: 6JPI) (c) BusR-promoter DNA complex (PDB ID: 7B5Y). Coarse-grained representations of the DNA-protein complexes are shown below the scattering profiles. In the snapshots, the DNA molecule is rendered in cyan, and the protein subunits are in different colors using the space-filling representation. For each complex, the simulations were carried out at different ionic strengths (0.5 M for (a), 0.3 M for (b) and 0.1 M for (c)) to mimic the solution conditions in the scattering experiments. In (a)-(c), 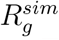 and 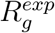 denote the radius of gyration estimated from simulations, and from a Guinier analysis of the experimental SAXS profiles, respectively. For all three complexes, these values are in excellent agreement with experiments.

### Comparison with experimental chemical shifts

In contrast to SAXS, which is useful for elucidating the global dimensions of biomolecules, NMR provides structural and dynamical information at the atomic scale.^102^ To evaluate if COFFEE faithfully captures the conformational fluctuations within a NMR-derived structural ensemble. We considered the homodimeric Lac Repressor DNA binding domain (DBD) bound to its natural operator, O1 DNA.^101^ In this complex, helix-turn-helix (HTH) motifs within the DBD bind specifically to the major groove of the operator DNA. A small residue fragment adjacent to the DBD, known as the helical hinge domain (residues *≈* 50-58) inserts into the DNA minor groove, lending additional stability. Given its small size, this complex has been extensively studied using molecular dynamics simulations^37,103–105^ to infer details of DNA-protein binding specificity in the context of transcription regulation.

We initiated simulations at 315 K using the coordinates of the NMR-derived structure deposited in the PDB database (PDB ID: 1L1M).^101^ The residue-wise C_*α*_ chemical shifts for the two independent subunits of the DBD domain determined using the LARMOR-C_*α*_ formalism are shown in Fig. 5(a) and 5(b). As is evident, our simulations accurately repro-duce the experimental chemical shifts. ^101^ However, we do observe *≈* 2-3 ppm downshift for some residues within the HTH motifs, indicating a marginal loss of *α*-helical character along the trajectory. It is important to note that the DNA-protein contact potential was calibrated to reproduce the B-factors reported for a low-temperature crystal structure (Fig. 2), and without any further readjustments to the force-field parameters, we obtain quantitative agreement with NMR chemical shifts (recorded at 315 K) for a completely unrelated DNA-protein complex.

**Figure 5:**
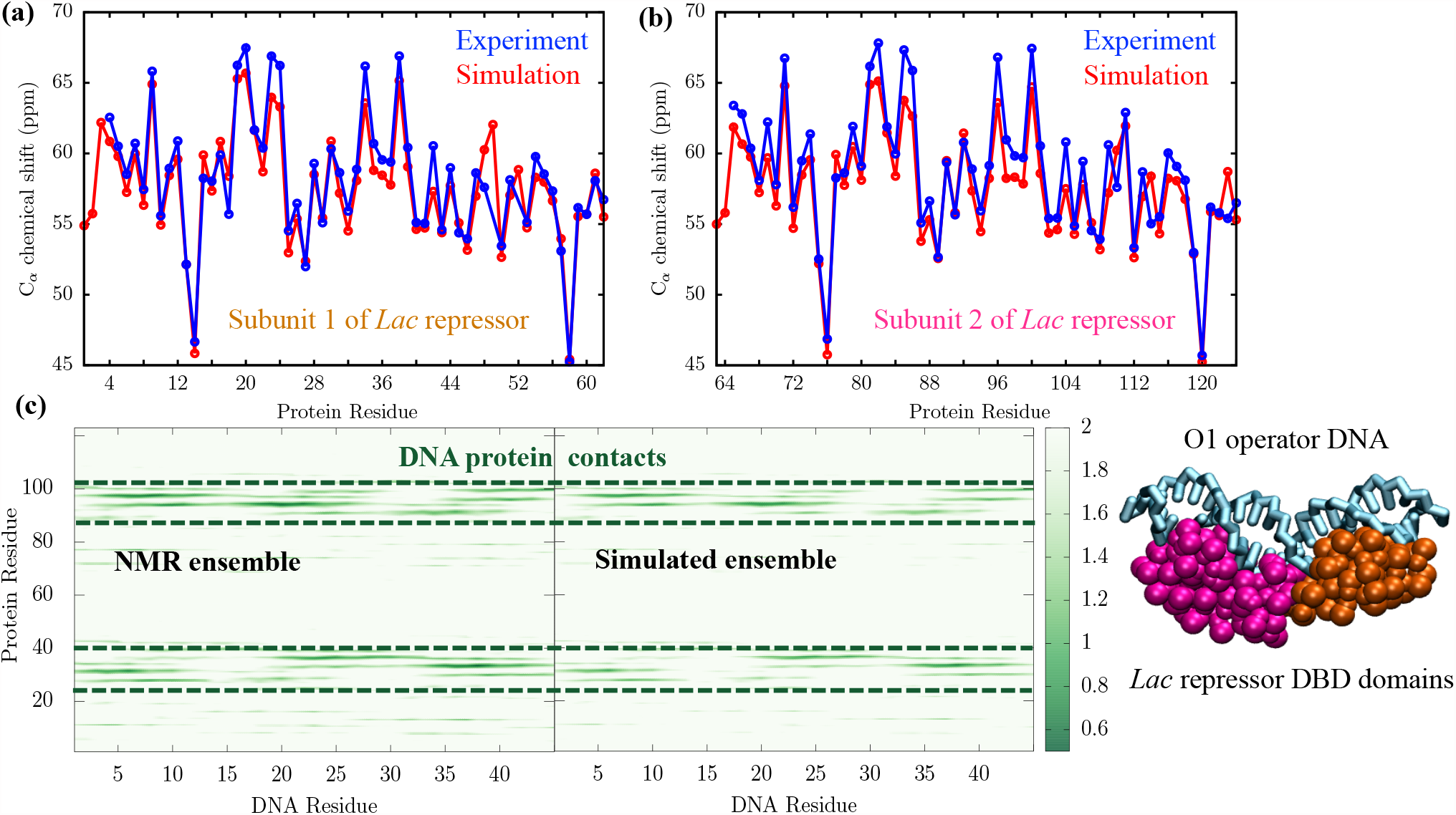
(a) and (b) Comparison of the experimental C_*α*_ chemical shifts (blue open circles)^101^ with those computed from simulations using the LARMOR-C_*α*_ formalism (red filled circles) for the two subunits of Lac repressor-DNA binding domain (DBD) when bound to its natural operator, O1. (c) DNA-protein contact maps computed from the NMR ensemble (left) and simulations (right). Intense colors in certain regions of the contact map (demarcated by dashed lines) indicate that residues are in close contact. A coarse-grained representation of the DNA-protein complex (PDB ID: 1L1M) is also shown. The O1 operator DNA rendered in cyan. The Lac repressor is rendered using a space-filling representation, with Subunit 1 in orange, and Subunit 2 in magenta.

The binding of the Lac Repressor to O1 is asymmetric, with the individual DBD subunits adopting different orientations within the complex. ^101^ As a result, the pattern of DNA-protein contacts established by the two subunits are distinct. The observed asymmetry is visible from the DNA-protein interaction maps shown in Fig. 5(c), particularly within the dashed lines, which demarcate the interactions of HTH motif within each DBD with the major grooves of the operator. In both the NMR and the simulated interaction maps (Fig. 5(c)), the intensity of certain pixels (which denote the contact distance between a DNA and a protein residue) is clearly different for equivalent positions on the two major grooves. Strikingly, the simulated ensemble retains the key residue-residue contacts resolved using NMR. The reduced pixel intensity in certain regions of the map, however, imply that some DNA-protein interactions (present in the NMR ensemble) are transiently broken along the simulation trajectory. This is not entirely unexpected given the dynamic nature of the Lac-DNA complex, and its ability to exploit multiple binding modes. ^103^

In the previous sections, we illustrated how COFFEE accurately reproduces different experimental observables, such as crystallographic B-factors, SAXS profiles, and C_*α*_ chemical shifts for diverse DNA-protein complexes. Of course, in these examples, the conformational ensembles are rather restricted and primarily include small fluctuations around the native basin. To test whether our model can describe large-scale conformational transitions with the same level of accuracy, we probe the salt-dependent unwrapping in nucleosome. These results are described in the following sections.

### COFFEE reproduces the experimental scattering curve for the nucleosome

In eukaryotic cells, genomic DNA is highly compacted in a hierarchical fashion, and packaged into a micron sized nucleus. At the nanometer scale, packaging occurs through the formation of nucleoprotein complex known as chromatin. The nucleosome core particle (or simply the nucleosome) is the basic repeating unit. In a nucleosome, *≈* 147 bp of double-stranded DNA wraps around an octameric core of histone proteins to form a left-handed superhelix consisting of *∼* 1.65 turns (Fig. 6(a)).^106,108^ A canonical histone core consists of a H3-H4 tetramer, and two H2A-H2B dimers. Each histone protein consists of three *α*-helices connected by intervening loops. ^106,108^ The N-termini of each histone consists of disordered tails, while H2A also has a tail at its C-terminus. The histone tails are hotspots for post-translational modifications (PTMs), and often mediate inter-nucleosome interactions within chromatin.^108^

**Figure 6:**
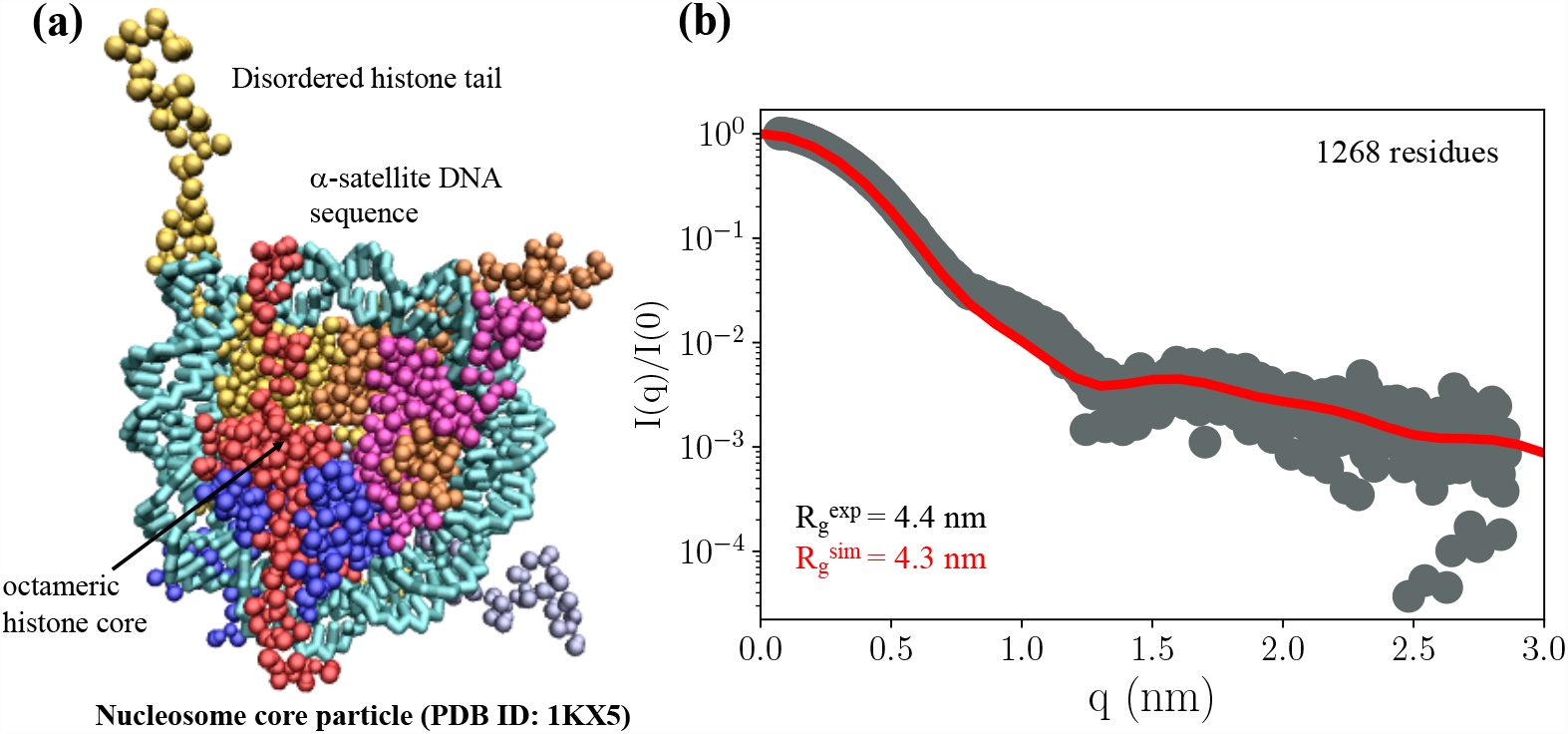
(a) A coarse-grained representation of the nucleosome core particle consisting of the *α*-satellite DNA sequence (PDB ID: 1KX5).^106^ DNA is rendered in cyan, and the histone proteins, including the disordered tails, are shown in different colors using a space-filling representation. (b) Structure factor calculated from simulations (red) and SAXS experiments (gray points) at a monovalent salt concentration of 0.2 M. The experimental profile^107^ corresponds to the 601 Widom sequence, which has the same number of basepairs, but a different composition than the *α*-satellite DNA sequence considered here. The 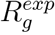 and 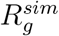 are the radii of gyration calculated from simulations, and from a Guinier analysis of the experimental SAXS profiles, respectively.

We simulated a nucleosome core particle consisting of 147 bp of palindromic dsDNA derived from the human *α*-satellite sequence repeat. The DNA helix is wrapped around *Xenopus laevis* core histones. The initial coordinates for this nucleosome sequence were taken from a high resolution crystal structure reported by Richmond and coworkers (Fig. 6(a)). ^106^ We compare the simulated scattering curve at 0.2 M with the experimental SAXS profile corresponding to a nucleosome reconstituted from the Widom 601 sequence. ^107^ Although the sequence composition of the *α*-satellite sequence differs greatly from Widom 601, the simulated scattering curve is almost superimposable on the experimental profile (Fig. 6(b)). Below *q ≈* 0.8 nm (the Guinier regime) the agreement between the simulated and experimental profiles is particularly impressive (Fig. 6(b)). In addition, the predicted radius of gyration 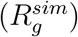 almost coincides with the experimental estimate 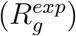, despite the differ-ences in the DNA sequence. The minor deviations at *q ≈* 0.8-1.0 nm could stem from the weaker positioning of the *α*-satellite DNA sequence (compared to the Widom 601 construct), which increases deformability at intermediate length-scales. Our observation suggests that at low salt concentration, where the nucleosome remains mostly in the wrapped configuration, the global dimension (as determined by *R*_*g*_) is insensitive to the variations in the DNA sequence. Sequence-specific features probably manifest themselves at shorter length-scales.

### Exploring salt-dependent nucleosome unwrapping using COFFEE

To probe nucleosome stability at different monovalent salt concentrations, we calculated *P*(*ϕ*_*UBP*_), the distributions of the number of unwrapped basepairs (see Eq. (11)-(13) for details). At *≈* 0.1 M, *P*(*ϕ*_*UBP*_) is extremely narrow and is centered around zero (Fig. 7), suggesting that the nucleosome prefers to be in the fully wrapped conformation, exhibiting only local conformational fluctuations. As the salt concentration is increased, electrostatic interactions between DNA and the histone core are weakened, and the population shifts towards partially unwrapped states. For instance, at *≈* 0.5 M, about 25 bp are detached from the histone core (Fig. 7)). The distributions also become progressively broader, suggesting that a wide array of conformations are accessible. At intermediate salt concentrations, *P*(*ϕ*_*UBP*_) curves feature multiple peaks. These correspond to metastable states exhibiting different extent of unwrapping. We find that the ensemble of partially unwrapped states is diverse, characterized by both asymmetric (snapshot A, Fig. 7), as well as symmetric detachment (snapshot S, Fig. 7) of DNA from the histone core. Even at a very high salt concentration (*≈* 1.0 M), DNA does not fully detach from the histone core. Indeed, in some trajectories, the free DNA partially folds back and interacts with the disordered histone tails. There is no evidence of histone loss, although experiments^57^ suggest that H2A-H2B dimers may dissociate at *≈* 1-1.5 M, leading to the formation of sub-nucleosomal particles.

**Figure 7:**
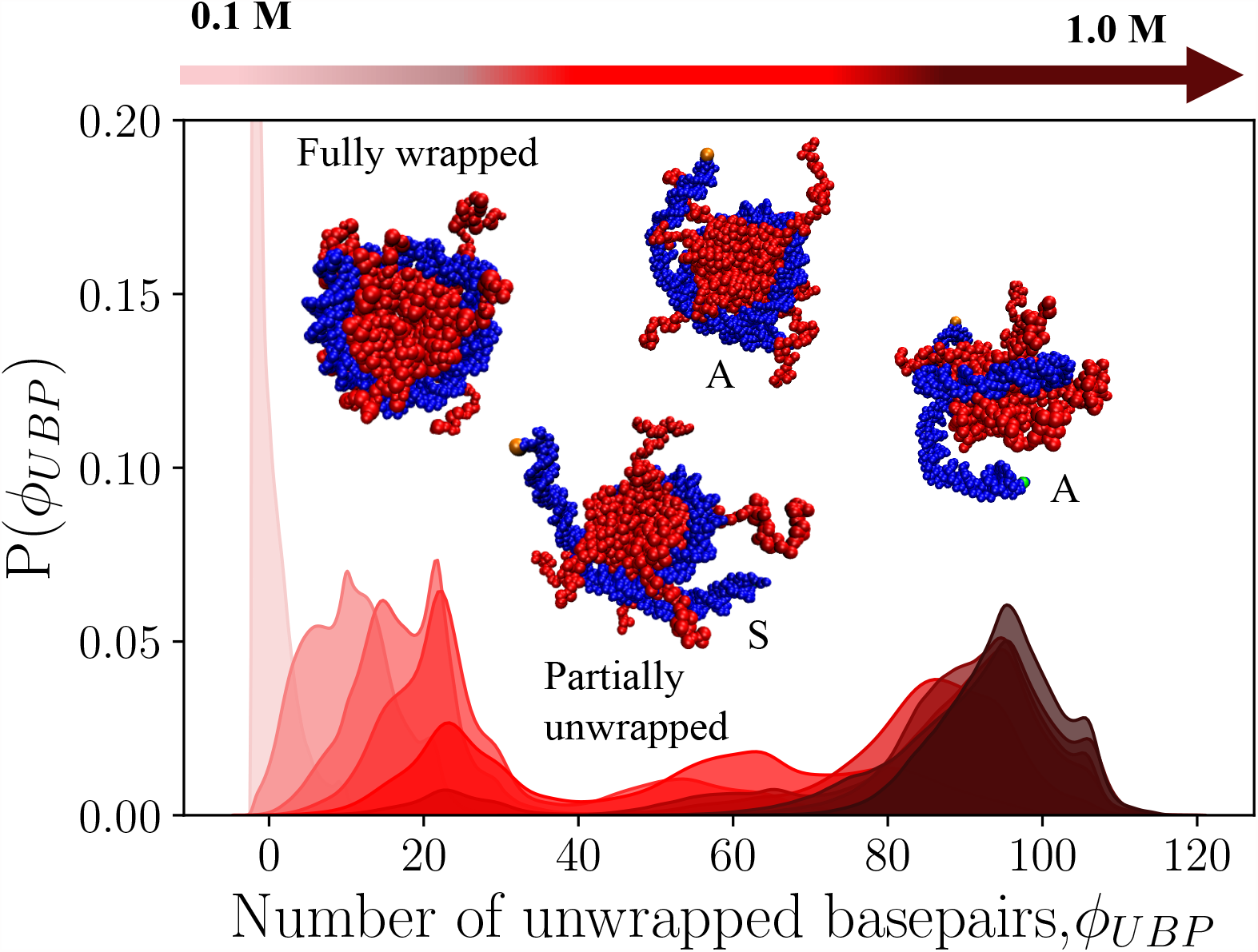
The probability distributions of *ϕ*_*UBP*_, the number of unwrapped base-pairs at different salt concentrations. The nucleosome core particle consists of the human *α*-satellite DNA sequence (shown in blue) wrapped around an octamer of *Xenopus laevis* histones (shown in red). At low salt-concentrations, the nucleosome is in the fully wrapped configuration. Various partially unwrapped states (asymmetric (A), and symmetric (S)) are populated as the salt concentration is increased. Some representative snapshots are shown superimposed on the graph.

The stabilization of different unwrapped states with increase in salt concentration suggests that nucleosome disassembly is likely to proceed via multiple pathways, with the finer details of the mechanism being dependent on the sequence as well as other external factors. Recent SAXS experiments, as well as ensemble optimization techniques,^56,57^ which generate pools of structures that are compatible with experimental data, already hint at such a scenario.

### Deletion of histone tails destabilizes the nucleosome

Within the nucleosome, the disordered histone tails often function as gatekeepers, preventing unwarranted sliding or unwrapping. They are rich in positively charged amino acids (LYS and ARG), which form specific interactions with the negatively charged phosphate backbone. Truncation or deletion of histone tails destablize the nucleosome,^109,110^ affecting not only nucleosome repositioning, but also higher order chromatin organization.

The effects of histone tail deletion on nucleosome stability are accurately captured by our model. As shown in Fig. 8(a), at low salt concentrations (*≈* 0.1 M), the tailless nucleosome preferentially populates partially unwrapped states. The average number of unwrapped base pairs, ⟨ *ϕ*_*UBP*_ ⟩, is also consistently higher at all salt concentrations for the tailless nucleosome (Fig. 8(b)). We find that the *ϕ*_*UBP*_ distributions at intermediate salt concentrations are typically broader than those depicted in Fig. 7. This trend implies that the tailless nucleosome is indeed more pliant, and can readily switch between alternate conformations. The unwrapped states exhibit both symmetric, as well as asymmetric DNA detachments (snapshots A and S, Fig. 8(a)). Interestingly, at *≈* 1.0,M, we observe a small peak in *P*(*ϕ*_*UBP*_), corresponding to the fully unwrapped state (snapshot U, Fig. 8(a)). Even after completely disassembly, the histone core remains completely intact, consistent with the pioneering studies of Kornberg ^111^ and Moudrianakis, ^112^ which suggest that the octamer readily breaks down only at low salt concentrations.

**Figure 8:**
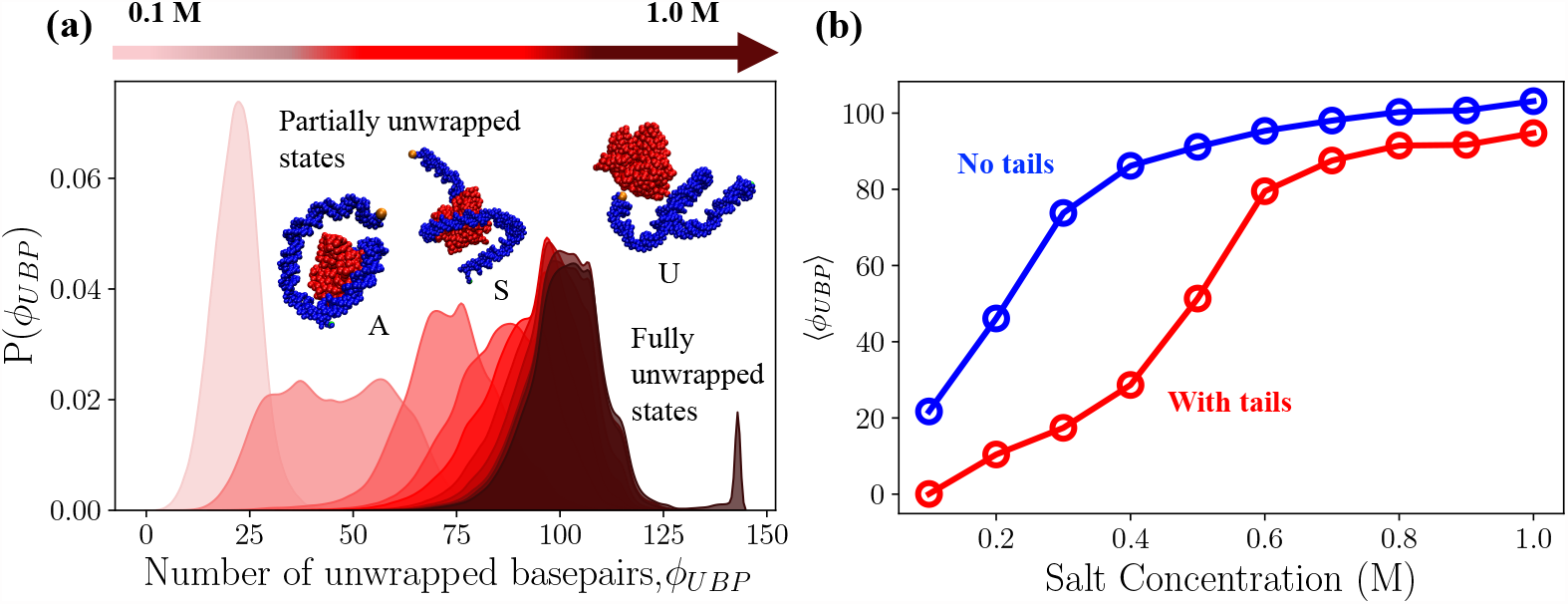
The effect of tail removal for a nucleosome consisting of 147 bp of human *α*-satellite DNA wrapped around an octameric core of *Xenopus laveis* histones. (a) The probability distributions of *ϕ*_*UBP*_, the number of unwrapped base-pairs at different salt concentrations. DNA is rendered in blue, and the histone core in red. (b) The variation of the average number of unwrapped base-pairs, ⟨ *ϕ*_*UBP*_ ⟩, with salt concentration for a nucleosome with tails (red) and without tails (blue).

### Mutations at superhelical locations enhance nucleosome flexibility

Besides electrostatic complementarity, the nucleosome complex is also stabilized by non-covalent interactions of chemical nature, which provide additional finesse during genome organization.^106^ Among these interactions, Arginine-phosphate salt-bridges at specific superhelical locations where the nucleosomal DNA makes contacts with the histone core, are the most critical.^113,114^ Indeed, disruption of these ‘special’ contacts are associated with SIN mutations in yeast, and are known to enhance nucleosome accessibility.^115^

To probe if COFFEE is sensitive to chemical perturbations, we simulated a mutant nucle-osome sequence, where the Arginines at eight superhelical locations are replaced with Lysines (Fig. 9(a)). Both ARG and LYS have a charge of +1 at neutral pH, and hence changes in stability cannot be explained by electrostatic interactions alone. The *ϕ*_*UBP*_ distributions at M for the canonical and the mutant nucleosome are shown in Fig. 9(b). Both sequences exhibit a bimodal distribution, suggesting the presence of at least two metastable states. For the canonical nucleosome, a partially unwrapped structure with *ϕ*_*UBP*_ *≈* 30 is the major state, while structures exhibiting more extensive unwrapping (*ϕ*_*UBP*_ *≈* 60) are less populated. As is evident from Fig. 9(b), the population shifts in favor of highly unwrapped states (*ϕ*_*UBP*_ *≈* 80) in the ARG*→*LYS mutant nucleosome, with partially unwrapped configurations (*ϕ*_*UBP*_ *≈* 30, similar to those found in the canonical nucleosome) being substantially destabilized. Hence, mutating ARG to LYS substantially weakens DNA-protein contacts and enhances nucleosome flexibility. This effect becomes particularly important in the context of CENP-A nucleosomes, in which ARG to LYS replacement facilitates proteolysis, thereby preventing promiscuous assembly, and providing a clearance mechanism from euchromatin regions.^58,59^ The ARG to LYS mutation clearly does not alter the net charge of the nucleosome complex, which implies that there is at best only a minor change in the electrostatic interactions. To explain the reduced stability in the LYS mutant requires accounting for specific chemical effects, which are not considered in many CG models. In instances where sequence specificity is important, brewing COFFEE would be ideal method.

**Figure 9:**
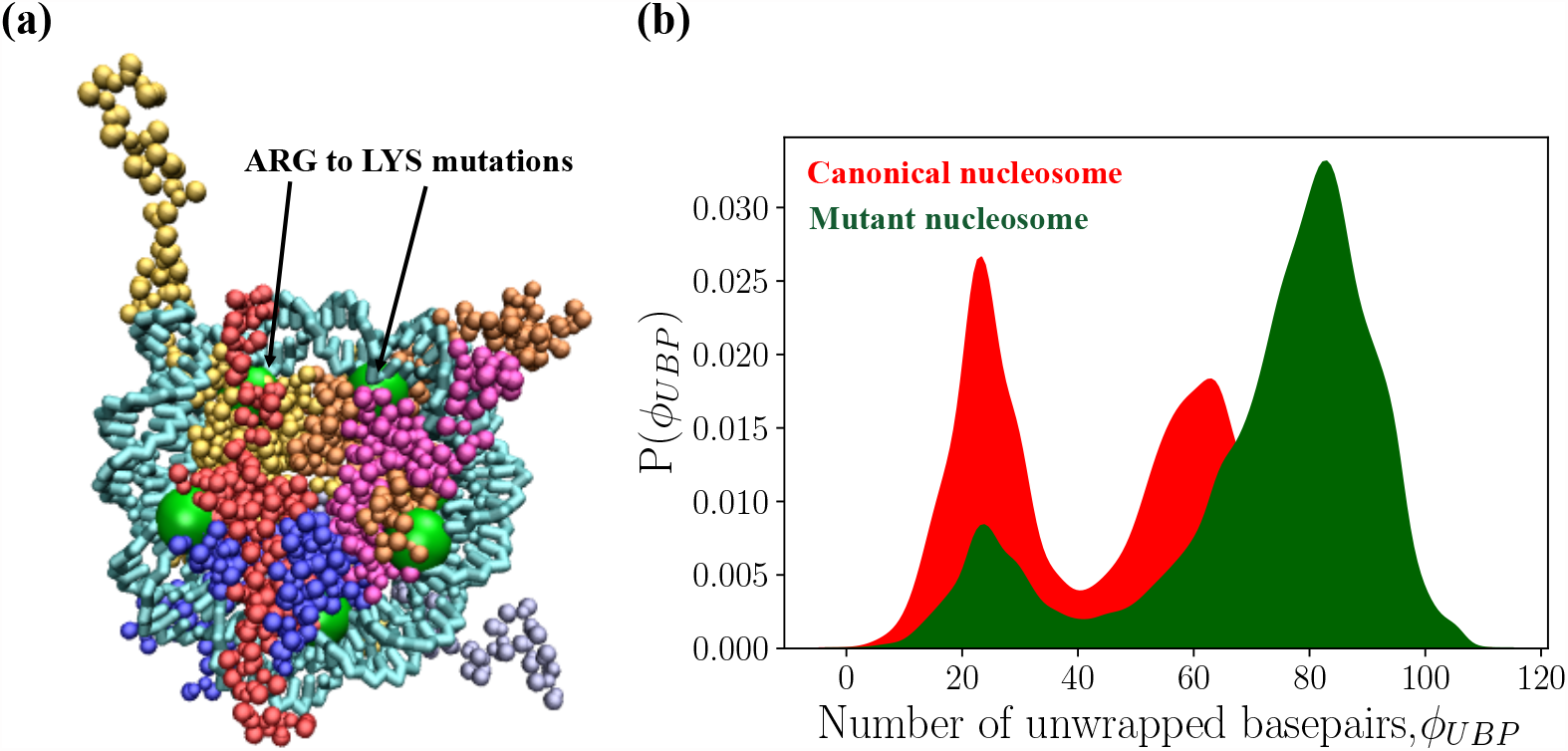
(a) A coarse-grained representation of the mutant nucleosome. The coloring scheme is the same as in Fig. 6(a). The green spheres denote the superhelical locations where Arginines are replaced by Lysines. (b) Probability distributions of *ϕ*_*UBP*_, the number of unwrapped base-pairs at a salt concentration of 0.5 M, for the canonical (red) and the mutant nucleosome (green).

## Concluding Remarks

In order to move towards a quantitative description of DNA-protein complexes, which are major components of the cellular machinery, we developed, COFFEE, a transferable coarse-grained (CG) model. Simulations based on COFFEE for a number of DNA-protein complexes demonstrate that it is a robust computational framework that takes into account the sequence-specific chemistry explicitly. The novel feature of COFFEE is that it de-scribes DNA-protein binding using statistical potential (SP) derived from a database of high-resolution structures. Incorporation of the SP into the previously introduced TIS model for DNA^53^ and the SOP-SC model for proteins^42^ results in COFFEE having a only single adjustable parameter (see below). Applications to a variety of DNA-protein complexes, including the nucleosome, attests to the accuracy and transferability of COFFEE.

### Force-field parameters and calibration

To brew COFFEE, we use the TIS model for DNA^53^ and the SOP-SC potential^39,60^ for folded proteins as the key ingredients. The TIS-DNA model,^53^ developed using a ‘top-down’ approach, includes sequence-dependent base-pairing and base-stacking interactions, which were calibrated to reproduce the thermodynamics of hairpin melting, and dimer stacking. The electrostatic interactions are described implicitly using a Debye-Hückel potential. Importantly, the TIS-DNA quantitatively reproduce many experimentally determined observables, as shown previously.^53^ The SOP-SC model for folded proteins, based on a similar conceptual framework, has been used with considerable success in studying protein folding in the presence of denaturants and pH. ^40,41,43^ A key ingredient of the SOP-SC energy function is a knowledge-based potential, ^72^ which encodes the sequence specific interaction between amino acids. These unique features of the TIS-DNA and SOP-SC force-fields, make them ideal for integration into COFFEE.

The TIS-DNA model has two adjustable parameters, which modulate the strength of the base-stacking and hydrogen-bonding interactions. The SOP-SC model has three adjustable energy scales that were adjusted to reproduce the melting temperatures of globular proteins.^39,42^ Here, we used the previously determined optimal values for the TIS-DNA,^53^ which reproduce the sequence-dependent melting temperatures of DNA hairpins. For the SOP-SC force-field, we adopted the parameter set that was used to predict the effect of pH on the folding thermodynamics and kinetics of ubiquitin. ^42^ We emphasize that TIS-DNA and SOP-SC have been combined in a modular fashion (“as is”), without requiring any re-optimization of the individual force-fields.

COFFEE is calibrated by adjusting a single parameter (*λ*_*DNAPRO*_), describing the strength of the non-covalent DNA-protein interactions. Strikingly, without any additional adjustments to the other force-field parameters, COFFEE faithfully reproduces the crystallographic B-factors for DNA-protein complexes of diverse shapes and sizes. Furthermore, COFFEE reproduces quantitatively the scattering profiles, as well as chemical shifts in quantitative agreement with experiments. The accuracy of COFFEE with (*λ*_*DNAPRO*_ being the only parameter) makes it an attractive transferable model for simulating arbitrary DNA-protein complexes.

### Conformational ensembles of nucleosomes are consistent with experiments

As a key application of COFFEE, we probed the salt-dependent conformational changes of a nucleosome core particle consisting of the *α*-satellite DNA sequence wrapped around *Xenopus laevis* core histones. The simulations *quantitatively* reproduce the dimensions and the scattering curve reported at 0.2 M. ^107^ We also show that diverse metastable states, exhibiting different extent of DNA detachment, are populated during salt-induced unwrapping, in accord with recent findings in time-resolved SAXS experiments.^57^ Interestingly, the nucleosome becomes more flexible when Arginines at certain super-helical locations are mutated to Lysines, which does not alter the electrostatic interactions. Such destabilization arise due to subtle chemical effects, which are unlikely to be captured without explicitly taking sequence effects into account. The changes in stability due to ARG*→*LYS mutations cannot be explained based on electrostatic interactions alone. These subtle effects are accurately described by the SP developed from contact statistics, which is an integral part of COFFEE.

### Resolving sequence effects in nucleosomes with COFFEE

DNA and protein sequences dictate the conformational dynamics of nucleosomes,^116–120^ as well as higher-order chromatin organization.^121^ A remarkable experiment by Ha and coworkers, ^120^ combining optical tweezers and FRET, brought the role of sequence into the spotlight. The authors showed that the unwrapping direction could be controlled by systematically varying the local DNA sequence within the outer and inner wraps of a nucleosome. In a biological context, sequence-encoded plasticity is important for fulfilling key regulatory roles. ^116^ For instance, strongly positioned sequences (akin to the Widom 601 sequence) are particularly enriched near intergenic and coding regions, where maintaining genome integrity is critical. On the other hand, weaker sequences, which unwrap easily, are abundant in highly transcribed regions.^116^

Despite these important insights, a microscopic picture of how sequence-encoded interactions predispose the nucleosome for spontaneous gaping/unwrapping, or invasion by chromatin remodellers is missing. The dust has also not completely settled on unidirectional unwrapping. ^120^ Whether it represents an universally preferred mode of DNA detachment in nucleosomes continues to be debated.^122^ Given that COFFEE blends robust models for describing sequence-dependent properties of DNA and proteins with a knowledge-based statistical potential for DNA-protein interactions, we anticipate that it would be a suitable framework for addressing these critical questions.

## Supporting Information Available

A description the SOP-SC and TIS-DNA force-fields; derivation of coarse-grained form factors using the isolated bead approximation (IBA); tabulated values of the SOP-SC and TIS-DNA force-field parameters; PDB IDs of complexes used for deriving the statistical potential (SP) for native DNA-protein contacts; Matrices denoting the effective energy scales for DNA-protein contacts; PDB IDs of DNA sequences used for deriving coarse-grained form factors; tabulated values of the coarse-grained form factors. Scripts and relevant input files for carrying out nucleosome simulations using the COFFEE framework within OpenMM are available from https://github.com/balaka92/Nucleosome.

This material is available free of charge via the Internet at http://pubs.acs.org/.

## Acknowledgement

We are grateful to Hung T. Nguyen and Sucheol Shin for fruitful discussions. We acknowledge the Texas Advanced Computing Center (TACC) for providing the necessary computing resources. Our work was supported by grants from the National Institutes of Health (GM-107703), National Science Foundation (CHE 2320256), as well as a grant from the Welch Foundation (F-0019) administered through the Collie-Welch Regents Chair.

The authors declare no competing interests.

