## Supporting Information for "Brewing COFFEE: A sequence-specific coarse-grained energy function for simulations of DNA-protein complexes"

**Self Organizing Polymer with Side-chains (SOP-SC) model for proteins:** The SOP-SC model[1,2] provides a quantitatively accurate description of the thermodynamics and kinetics of protein folding at different denaturant and salt concentrations, as well as pH.[3–7] In the SOP-SC model, each amino-acid is represented using two interaction sites: a backbone bead (BB) centered on the $C_\alpha$ atom and a side-chain bead (SC) placed on the center-of-mass of the side-chain (Figure S1). The energy function for the SOP-SC model is given by:

$$U_{SOP-SC} = U_{FENE} + U_N + U_{NN} + U_{ELE}^{PRO}. \tag{1}$$

The FENE potential, $U_{FENE}$, describing the chain connectivity of the protein molecule, is given by:

$$U_{FENE} = -\sum_{i=1}^{N_B^{PRO}} \frac{k}{2} R_0^2 \log\left(1 - \frac{(r_i - r_{i,ref})^2}{R_0^2}\right), \tag{2}$$

where $N_B^{PRO}$ denotes the total number of bonds present between the covalently linked beads, $r_i$ denotes the distance between $i^{th}$ pair of beads, and $r_{i,ref}$ is the equilibrium distance between the $i^{th}$ pair in the reference structure.

The non-bonded interactions are classified as either native ($U_N$) or non-native ($U_{NN}$). Interactions between two beads, which are at least separated by three bonds, are considered native, if the distance between them is less then a pre-defined cut-off, $R_c$. The native interactions, $U_N$, stabilizing the folded state, are described using a Lennard-Jones type potential given by:

$$\begin{aligned} U_N = &\sum_{i=1}^{N_N^{BB}} \epsilon_{BB}\left[\left(\frac{r_{ref,i}}{r_i}\right)^{12} - 2\left(\frac{r_{ref,i}}{r_i}\right)^{6}\right] + \sum_{i=1}^{N_N^{BS}} \epsilon_{BS}\left[\left(\frac{r_{ref,i}}{r_i}\right)^{12} - 2\left(\frac{r_{ref,i}}{r_i}\right)^{6}\right] + \\ &\sum_{i=1}^{N_N^{SS}} \epsilon_{SS}|\epsilon_i - 0.7|\left[\left(\frac{r_{ref,i}}{r_i}\right)^{12} - 2\left(\frac{r_{ref,i}}{r_i}\right)^{6}\right], \end{aligned} \tag{3}$$

In Eq. (3), $N_N^{BB}$, $N_N^{BS}$ and $N_N^{SS}$ are, the total number of native backbone-backbone, backbone-side-chain, and side-chain-side-chain interactions, respectively; $r_i$ is the distance between the $i^{th}$ pair of beads, and $r_{ref,i}$ is the distance between the same pair in the reference structure.

The three adjustable energy scales, $\epsilon_{BB}$, $\epsilon_{BS}$ and $\epsilon_{SS}$ are the strengths of the native interactions. The parameter $\epsilon_i$ denotes the Betancourt-Thirumalai matrix element,,[8] which encodes the sequence-specificity of the SOP-SC model.
The non-native interactions, $U_{NN}$, assumed to be purely repulsive, are modeled as:

$$U_{NN} = \sum_{i=1}^{N_{NN}} \epsilon_l \left(\frac{\sigma_i}{r_i}\right)^6 + \sum_{i=1}^{N_{NN}^{BB}} \epsilon_l \left(\frac{\sigma_{BB}}{r_i}\right)^6 + \sum_{i=1}^{N_{NN}^{BS}} \epsilon_l \left(\frac{\sigma_{BS}}{r_i}\right)^6, \tag{4}$$

In Eq. (4), $N_{NN}$ denotes the total number of non-native interactions between the protein beads; $N_{NN}^{BB}$ and $N_{NN}^{BS}$ denote the total number of angular interactions among backbone beads, and backbone-side-chain beads, which are separated by two bonds; $\sigma_i$ is the sum of the radii of the beads involved in the $i^{th}$ pair of interactions, $\sigma_{BB}$ denotes the diameter of the backbone bead, and $\sigma_i^{BS} = \beta\left[\sigma_{BB} + \sigma_{SC}\right]$ is the sum of the radii of the backbone and side-chain beads involved in the $i^{th}$ pair of angular interactions, with $\beta$ denoting a scaling factor. Following earlier work,[9,10] we set $\beta = 0.8$.

The electrostatic interactions involving the charged amino-acid side-chains are described using a Debye-Hückel potential.

$$U_{ELE}^{PRO} = \sum_{i,j}{}' \frac{q_i^{PRO} q_j^{PRO} \exp\left(-\kappa r_{ij}\right)}{\epsilon_{PRO} r_{ij}}, \tag{5}$$

where $q_i^{PRO}$ is either $+1e$ or $-1e$ depending on the amino-acid side-chain, $\kappa^{-1}$ is the inverse Debye length, and $r_{ij}$ denotes the distance between the side-chains of charged residues $i$ and $j$. Following earlier work,[9] the dielectric constant, $\epsilon_{PRO}$ is set to 10.0. The values of the different parameters of the SOP-SC model are listed in Tables S1 and S2.

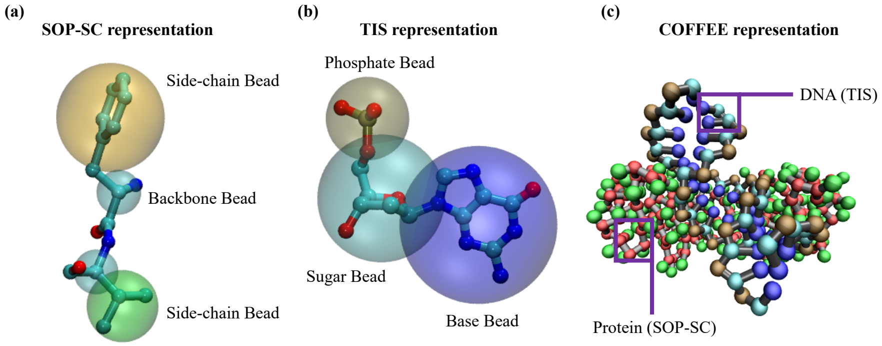


**Figure S1.** (a) A schematic illustration of the coarse-grained representation used in the SOP-SC model for proteins. Each amino-acid is represented by two beads. One bead is centered on the backbone $C_\alpha$ atom, and the other one is centered on the center-of-mass of the side-chain. The side-chain beads are shown in different colors to illustrate the sequence-specificity encoded in the SOP-SC model. (b) The coarse-grained representation used in the TIS model of DNA. Each nucleotide is represented using three spherical beads, positioned on the centers of mass of the phosphate, the sugar group, and the base. (c) COFFEE representation of the Human TBP core-domain complexed with DNA (PDB ID: 1CDW), in which the DNA and the protein chains are coarse-grained to the TIS and SOP-SC resolutions, respectively.

**The Three Interaction Site (TIS) model for DNA:** In the TIS model for DNA,[11] each nucleotide is represented by three beads, which are positioned on the centers of mass of the phosphate, sugar, and the base (Figure S1). It provides a remarkably accurate description of the sequence-dependent properties of single-stranded and double-stranded DNA.[11] The total energy, $U_{TIS}$, for a given conformation of the polynucleotide includes contributions from the bond, angular, stacking, hydrogen-bonding, excluded-volume, and electrostatic interactions:

$$U_{TIS} = U_B + U_A + U_S + U_{HB} + U_{EV} + U_{ELE}^{DNA}. \tag{6}$$

The bond and angular interactions are described using harmonic potentials.

$$U_B = k_r(r - r_{ref})^2 \tag{7}$$

$$U_A = k_\alpha(\alpha - \alpha_{ref})^2. \tag{8}$$

In Eq. (7) and (8), $r_{ref}$ and $\alpha_{ref}$ denote the equilibrium bond lengths and bond angles found in the reference structure, and $k_r$ and $k_\alpha$ denote the force constants. The values of $k_r$ and $k_\alpha$ for the different bonds and angles are tabulated in Tables S3 and S4, respectively.
The volume exclusions between the interaction sites were described using the Weeks-Chandler-Andersen potential:[12]

$$U_{EV} = \epsilon_0 \left[ \left( \frac{D_0}{r} \right)^{12} - 2 \left( \frac{D_0}{r} \right)^6 + 1 \right], r < D_0 \tag{9}$$

where $D_0 = 3.2$ Å and $\epsilon_0 = 1$ kcal/mol. To minimize the number of parameters without affecting the accuracy of the results, all the interaction sites were assigned the same values of $D_0$ and $\epsilon_0$. As discussed in the context of the TIS model for RNA,[13] this particular choice of $D_0$ modestly underestimates the distance of closest approach between the interaction sites, but does not affect the folding thermodynamics.
The stacking interaction between two consecutive nucleotides along the DNA chain is given by:

$$U_S = U_S^0(1 + k_l(l - l_{ref})^2 + k_\phi(\phi_1 - \phi_1^{ref})^2 + k_\phi(\phi_2 - \phi_2^{ref})^2)^{-1}. \tag{10}$$

The deviations from the equilibrium geometry from a reference structure (described by the stacking distance $l_{ref}$ and backbone dihedrals, $\phi_1^{ref}$, and $\phi_2^{ref}$) modulate the strength of the stacking interactions. The values of the force constants, $k_l$ and $k_\phi$ are given in Table S5. The geometric parameters ($l$, $\phi_1$ and $\phi_2$)), which describe the stacking interactions in the TIS model are shown in Fig. S2. In Eq. (10), $U_S^0$ describes the stacking interaction for a particular dimer. It was calibrated to reproduce the thermodynamics, as described by the nearest-neighbor model of SantaLucia and Hicks.[14,15] In terms of enthalpic and entropic contributions, the strength of the stacking interaction, $U_S^0$, can be expressed as:

$$U_S^0 = -h + k_B\,(T - T_m)\,s, \tag{11}$$

where $h$ and $s$ are adjustable parameters, $T_m$ is the melting temperature of a specific dimer. The optimal values of $h$ and $s$, and $T_m$ for each dimer are tabulated in Table S6.
The hydrogen-bonding interactions are only considered between the canonical (Watson-Crick) base-pairs. The potential, $U_{HB}$ is given by:

$$U_{HB} = \frac{U_{HB}^0}{1 + k_d(d - d_0)^2 + k_\theta(\theta_1 - \theta_1^0)^2 + k_\theta(\theta_2 - \theta_2^0)^2 + k_\psi(\psi_1 - \psi_1^0)^2 + k_\psi(\psi_2 - \psi_2^0)^2 + k_\psi(\psi_3 - \psi_3^0)^2}. \tag{12}$$

In Eq. (12), $U_{HB}^0$ denotes the strength of the hydrogen-bonding interaction. The equilibrium value of the hydrogen bond distance is $d$, and and $\theta_1$, $\theta_2$, $\psi_1$, $\psi_2$, and $\psi_3$ denote angles and dihedrals that modulate the strength of the interaction. In Figure S2, we show a graphical illustration of the structural parameters described in Eq. (12).

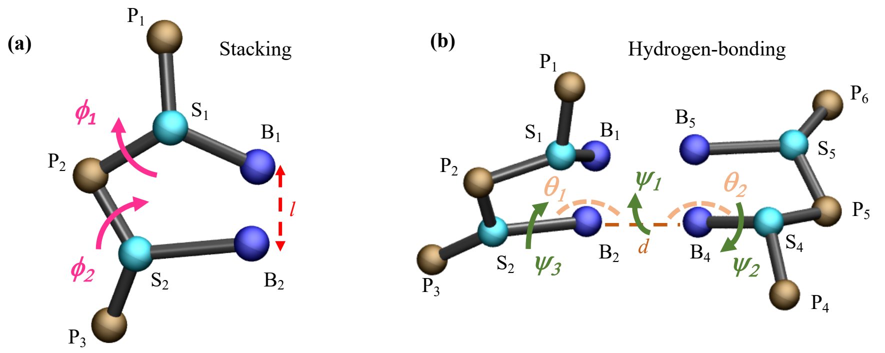


**Figure S2.** (a) Schematic of a dimer in the TIS representation, with $l$ ($B_1$–$B_2$) denoting the stacking distance, and $\phi_1$ ($P_1$–$S_1$–$P_2$–$S_2$) and $\phi_2$ ($P_3$–$S_2$–$P_2$–$S_1$) denoting the backbone dihedrals. (b) The hydrogen-bonding distance, $d$, between sites $B_2$ and $B_4$ is shown using a dotted line. The angles $\theta_1$ ($S_2$–$B_2$–$B_4$) and $\theta_2$ ($S_4$–$B_4$–$B_2$), and torsions $\psi_1$ ($S_2$–$B_2$–$B_4$–$S_4$), $\psi_2$ ($B_2$–$B_4$–$S_4$–$P_5$), and $\psi_3$ ($B_4$–$B_2$–$S_2$–$P_3$) which modulate the strength of the hydrogen-bonding interactions, are also labeled. Figure is adapted from our previous work.[11]

The electrostatic interactions between the phosphates are computed using the Debye-Hückel approximation, in conjunction with the Oosawa-Manning counterion condensation mechanism.[16] The electrostatic energy, $U_{ELE}^{DNA}$ is given by:[17]

$$U_{ELE}^{DNA} = \frac{q_{DNA}^2}{2\epsilon_{DNA}} \sum_{i,j} \frac{\exp(-\kappa r_{ij})}{r_{ij}}, \tag{13}$$

where $r_{ij}$ is the distance between two phosphates $i$, and $j$, $\epsilon_{DNA}$ is the dielectric constant of water, and $\kappa^{-1}$ is the Debye-screening length.

The magnitude of the phosphate charge, $q_{DNA}$ is determined using the Oosawa and Manning counterion condensation theory. The bare charge on the phosphate is renormalized due to the propensity of counterions to condense around DNA. The renormalized charged on the phosphate is,

$$q_{DNA} = q'_{DNA}(T) = \frac{b}{l_B(T)}, \qquad l_B(T) = \frac{e^2}{\epsilon k_B T}, \tag{14}$$

where $b$ is the length per unit charge, and $l_B$ is the Bjerrum length. The length per unit charge for DNA, as estimated by Olson and coworkers,[18] is approximately 4.4 Å, which leads to a reduced charge of $-0.6$ for the phosphates at 298 K.

**Derivation of the coarse-grained form factors:** For a molecule consisting of $N$ atoms, the scattering amplitude at vector $\mathbf{q}$ is given by:

$$A(\mathbf{q}) = \sum_{i=1}^{N} f_i^{aa}(q) \exp(i\mathbf{q}.r_i), \tag{15}$$

where $f_i^{aa}(q)$ and $r_i$ denote the scattering factor and position of atom $i$. If the molecules are assumed to be randomly oriented (as would be the case in solution), orientational averaging can be carried out, and the scattering intensity can be calculated using the Debye formula:

$$I(q) = \langle I(q) \rangle = \sum_{i=1}^{N} \sum_{j=1}^{N} f_i^{aa}(q) f_j^{aa}(q) \frac{\sin(qr_{ij})}{qr_{ij}}. \tag{16}$$

The atomic form factor, $f_i^{aa}(q)$ can be computed from the Cromer-Mann analytic function:[19,20]

$$f_i^{aa}(q) = \sum_{k=1}^{4} a_k \exp\left[-b_k \left(\frac{q}{4\pi}\right)^2\right] + c, \tag{17}$$

where the parameters $a_k$, $b_k$ and $c$ for each element are taken from the International Tables for Crystallography.[20] The form factors computed using Eq. (17) are only effective in estimating the scattering intensities in vacuum. Since the SAXS experiments are carried out in solution, additional corrections must be incorporated to provide a realistic description of the scattering profiles. To take into account the effect of the displaced solvent, an approximation proposed by Fraser *et al.*[21] is generally employed,

$$f_i'^{aa}(q) = f_i^{aa}(q) - \nu_i \rho_b \exp\left(\frac{-q^2 \nu_i^{2/3}}{4\pi}\right) \tag{18}$$

In Eq. (18), $\rho_b$ is the electron density of bulk water (taken to be 0.334 $e\mathring{A}^{-3}$), and $\nu_i$ is the van der Waals volume of atom $i$.[22] The modified form factors, $f_i'^{aa}(q)$, are used in Eq. (16) to provide a more realistic description of scattering in the solution state.
*Isolated Bead Approximation:* When a biomolecule is described using a coarse-grained representation, rather than in atomic detail, the scattering intensity is approximately given by:

$$I(q) \approx \sum_{i=1}^{m} \sum_{j=1}^{n} f_i(q) f_j(q) \frac{\sin(qR_{ij})}{qR_{ij}} \tag{19}$$

where $f_i(q)$ and $f_j(q)$ denote the coarse-grained form factors of beads $i$ and $j$, and $R_{ij}$ is the distance between them. Several approaches for computing $f(q)$ have been suggested,[22–24] including the isolated bead approximation (IBA),[24] which is simple, yet accurate. In the IBA, $f(q)$, is calculated as the square-root of the scattering intensity of all the atoms grouped within the coarse-grained bead, and then averaged over all the possible conformations in a dataset of experimental structures. In other words, $f(q)$ can be expressed as:

$$f(q) = \left\langle \left[\sum_{k \in i} \sum_{l \in i} f_k'^{aa}(q) f_l'^{aa}(q) \frac{\sin(qr_{kl})}{qr_{kl}}\right]^{1/2} \right\rangle_{PDB} \tag{20}$$

In Eq. (20), $f_k'^{aa}(q)$ and $f_l'^{aa}(q)$ denote the corrected form factors for the atoms that are part of the $i^{th}$ coarse-grained bead. The ensemble average is performed over all possible orientations of the atom group found within a database of experimental structures.

The residue-specific form factors for the protein backbone and the different side-chains, derived using the IBA, were available from Table S2 of Tong *et al.*[22] The nucleotide-specific form factors corresponding to the TIS representation were derived using Eqs (17)-(20) and are tabulated in Table S15. The experimental structures that were used for carrying out the ensemble-averaging in Eq. (20) are tabulated in Table S14.

Table S1: Parameters for the SOP-SC model. The energy scales, $\epsilon_{BB}$, $\epsilon_{BS}$ and $\epsilon_{SS}$ were optimized to reproduce the experimental melting profile of Ubiquitin.[9,10]

| Parameter | Value |
|---|---|
| $R_0$ | 2.0 Å |
| $R_c$ | 8.0 Å |
| $k$ | 20.0 kcal/(mol Å$^2$) |
| $\epsilon_l$ | 1.0 kcal/mol |
| $\epsilon_{BB}$ | 0.50 kcal/mol |
| $\epsilon_{BS}$ | 0.50 kcal/mol |
| $\epsilon_{SS}$ | 0.30 kcal/mol |

Table S2: Masses, Radii and charges of the coarse-grained beads used in the SOP-SC model.

| Residue | Mass (g/mol) | Radius (Å) | charge (e) |
|---|---|---|---|
| Gly | 1.00 | 0.50 | 0.0 |
| Ala | 12.01 | 2.52 | 0.0 |
| Val | 36.03 | 2.93 | 0.0 |
| Leu | 48.04 | 3.09 | 0.0 |
| Ile | 48.04 | 3.09 | 0.0 |
| Met | 68.09 | 3.09 | 0.0 |
| Phe | 84.07 | 3.18 | 0.0 |
| Pro | 36.03 | 2.78 | 0.0 |
| Ser | 28.01 | 2.59 | 0.0 |
| Thr | 40.02 | 2.81 | 0.0 |
| Asn | 54.02 | 2.84 | 0.0 |
| Gln | 66.04 | 3.01 | 0.0 |
| Tyr | 100.07 | 3.23 | 0.0 |
| Trp | 122.10 | 3.39 | 0.0 |
| Asp | 56.02 | 2.79 | -1.0 |
| Glu | 68.03 | 2.96 | -1.0 |
| His | 76.05 | 3.04 | 0.0 |
| Lys | 62.05 | 3.18 | 1.0 |
| Arg | 90.06 | 3.28 | 1.0 |
| Cys | 44.07 | 2.74 | 0.0 |
| backbone | 12.01 | 1.90 | 0.0 |

Table S3: The optimal values of the force constants, $k_r$, for the bond potentials in the TIS model.

| Bond Type | $k_r$(kcal/mol/Å$^2$) |
|---|---|
| SP | 62.59 |
| PS | 17.63 |
| SA | 44.31 |
| SG | 48.98 |
| SC | 43.25 |
| ST | 46.56 |

Table S4: The optimal values of the force constants, $k_\alpha$, for the angle-bending potentials in the TIS model.

| Angle Type | $k_\alpha$(kcal/mol/rad$^2$) |
|---|---|
| PSP | 25.67 |
| SPS | 67.50 |
| PSA | 29.53 |
| PST | 39.56 |
| PSG | 26.28 |
| PSC | 35.02 |
| ASP | 67.32 |
| TSP | 93.99 |
| GSP | 62.94 |
| CSP | 77.78 |

Table S5: Masses and excluded volume radii of the different DNA beads.

| Residue | Mass (g/mol) | Radius (Å) |
|---|---|---|
| DP | 94.95 | 2.0 |
| DS | 76.05 | 2.9 |
| DG | 146.08 | 3.0 |
| DA | 130.08 | 2.8 |
| DC | 106.06 | 2.7 |
| DT | 120.07 | 2.7 |

Table S6: The optimal values of the force constants for the stacking and hydrogen-bonding potentials in the TIS model of DNA.

| Parameter | Value |
|---|---|
| $k_l$ | 1.45Å$^{-2}$ |
| $k_\phi$ | 3.00 radians$^{-2}$ |
| $k_d$ | 4.00 Å$^{-2}$ |
| $k_\theta$ | 1.50 radians$^{-2}$ |
| $k_\psi$ | 0.15 radians$^{-2}$ |

Table S7: The parameters $h$, $s$ and $T_m$ for the different dimer stacks within the TIS model.

| $\frac{x}{w}$ | $h$ kcal/mol | $s$ | $T_m(K)$ |
|---|---|---|---|
| $\frac{A}{A}$ | 5.69 | 0.94 | 322.0 |
| $\frac{A}{T}$; $\frac{T}{A}$ | 4.95; 5.02 | 0.87; 0.65 | 293.0 |
| $\frac{A}{C}$; $\frac{C}{A}$ | 4.98; 5.03 | 0.92; 0.78 | 293.0 |
| $\frac{A}{G}$; $\frac{G}{A}$ | 5.43; 5.42 | 1.06; 0.98 | 333.6 |
| $\frac{C}{T}$; $\frac{T}{C}$ | 4.13; 4.14 | 0.71; 0.94 | 288.3 |
| $\frac{C}{C}$ | 4.12 | 0.92 | 288.3 |
| $\frac{C}{G}$; $\frac{G}{C}$ | 5.21; 5.13 | 2.31; 2.41 | 331.9 |
| $\frac{G}{T}$; $\frac{T}{G}$ | 5.28; 5.43 | 1.69; 1.72 | 332.6 |
| $\frac{G}{G}$ | 5.66 | -0.29 | 353.9 |
| $\frac{T}{T}$ | 4.17 | 0.89 | 288.3 |

Table S8: PDB IDs of complexes used for deriving the statistical potential (SP) for native DNA-protein contacts.

2NTC 2NP6 2NLL 2ISZ 2IS6 2IRF 2IIE 2IHM 2I13 2I06 2HHX 2HDD 2HAN 2H7H 2H7G 2H27 2GIH 2GE5 2G1P 2FQZ 2FLP 2FKC 2FCC 2EZV 2EX5 2E52 2E1C 2DTU 2DGC 2DDG 2D5V 2C9L 2C6Y 2BOP 2BNW 2B9S 2AQ4 2AOR 2AC0 2A66 2A07 1ZS4 1ZRF 1ZME 1YO5 1YF3 1Y6F 1XBR 1W7A 1W0U 1W0T 1V15 1UBD 1U8B 1TRR 1TKD 1TEZ 1TDZ 1TC3 1SKR 1SKN 1SFU 1SA3 1RXW 1RRS 1RM1 1RIO 1RH6 1R8E 1R7M 1R71 1R4O 1R0O 1QTM 1QRV 1QPZ 1QPI 1QNE 1QAJ 1PUF 1PUE 1PT3 1PP7 1PER 1PDN 1P71 1P47 1P3L 1OZJ 1ORN 1OE4 1NVP 1NLW 1NKP 1NH2 1NFK 1N48 1MUS 1MOW 1MNM 1MJ2 1MEY 1M5X 1M19 1M0E 1LQ1 1LMB 1LLM 1LE8 1L3L 1L2B 1KX5 1KU7 1KBU 1K82 1K79 1K78 1K61 1K4T 1K3W 1JJ4 1JGG 1JEY 1JE8 1J75 1J3E 1J1V 1IXY 1IHF 1IGN 1IG7 1IAW 1I3J 1HWT 1HU0 1HLV 1HCR 1HCQ 1H89 1H6F 1GXP 1GTW 1GD2 1GD2 1G9Y 1FYL 1FJL 1FIU 1F6O 1F4K 1F2I 1EYU 1EWQ 1ESG 1EQZ 1EMH 1EGW 1E3O 1DUX 1DSZ 1DP7 1DMU 1DIZ 1DC1 1D3U 1D2I 1D02 1CW0 1CKT 1CKQ 1CEZ 1BPX 1BL0 1BG1 1BDT 1B94 1B8I 1B8I 1B72 1B72 1B3T 1AZP 1AWC 1AU7 1AM9 1AKH 1A73 1A6Y 1A3Q 1A35 9ANT 3F21 3ERE 3E6C 3E54 3DPG 3CVU 3CRO 3COQ 3CO6 3CBB 3C2I 3C25 3C1B 3BTX 3BS1 3BRG 3BRF 3BRD 3BM3 3BIE 3BEP 2ZO1 2Z3X 2YVH 2VY1 2VS7 2VOA 2VLA 2VE9 2RBF 2RAM 2R9L 2R1J 2QOJ 2QL2 2QL2 2QHB 2PYJ 2PI0 2P0J 2OWO 2OFI 2ODI 2OAA 2O49

Table S9: Matrix denoting the effective energy scales (in kcal/mol) for DNA-protein contacts between the phosphate group (P) and amino-acid backbone (BB).

| | GUA | ADE | CYT | THY |
|---|---|---|---|---|
| ALA | 0.41 | 0.27 | 0.31 | 0.40 |
| GLY | 0.62 | 0.58 | 0.52 | 0.54 |
| LEU | 0.17 | 0.08 | 0.15 | 0.19 |
| ILE | 0.38 | 0.34 | 0.33 | 0.29 |
| TYR | 0.33 | 0.24 | 0.32 | 0.30 |
| TRP | 0.27 | 0.33 | 0.35 | 0.35 |
| LYS | 0.69 | 0.62 | 0.66 | 0.66 |
| PRO | 0.37 | 0.36 | 0.35 | 0.39 |
| GLU | 0.12 | 0.06 | 0.13 | 0.04 |
| HIS | 0.72 | 0.58 | 0.43 | 0.56 |
| ASP | 0.38 | 0.20 | 0.31 | 0.01 |
| ARG | 0.82 | 0.68 | 0.65 | 0.70 |
| GLN | 0.49 | 0.41 | 0.41 | 0.43 |
| ASN | 0.60 | 0.51 | 0.50 | 0.55 |
| CYS | 0.54 | 0.22 | 0.25 | 0.27 |
| SER | 0.70 | 0.56 | 0.59 | 0.60 |
| MET | 0.36 | 0.33 | 0.34 | 0.30 |
| VAL | 0.43 | 0.32 | 0.25 | 0.32 |
| THR | 0.67 | 0.57 | 0.57 | 0.64 |
| PHE | 0.44 | 0.25 | 0.26 | 0.33 |

Table S10: Matrix denoting the effective energy scales (in kcal/mol) for DNA-protein contacts between the sugar group (S) and amino-acid backbone (BB).

| | GUA | ADE | CYT | THY |
|---|---|---|---|---|
| ALA | 0.43 | 0.30 | 0.38 | 0.34 |
| GLY | 0.71 | 0.69 | 0.66 | 0.62 |
| LEU | 0.07 | 0.03 | 0.06 | 0.06 |
| ILE | 0.33 | 0.31 | 0.30 | 0.22 |
| TYR | 0.33 | 0.30 | 0.35 | 0.24 |
| TRP | 0.20 | 0.44 | 0.20 | 0.32 |
| LYS | 0.74 | 0.66 | 0.71 | 0.66 |
| PRO | 0.42 | 0.43 | 0.42 | 0.41 |
| GLU | 0.11 | 0.02 | 0.07 | 0.00 |
| HIS | 0.65 | 0.66 | 0.51 | 0.57 |
| ASP | 0.37 | 0.24 | 0.37 | 0.02 |
| ARG | 0.83 | 0.75 | 0.73 | 0.75 |
| GLN | 0.51 | 0.48 | 0.48 | 0.40 |
| ASN | 0.69 | 0.61 | 0.60 | 0.63 |
| CYS | 0.38 | 0.16 | 0.24 | 0.28 |
| SER | 0.85 | 0.63 | 0.74 | 0.66 |
| MET | 0.25 | 0.31 | 0.27 | 0.24 |
| VAL | 0.42 | 0.32 | 0.25 | 0.31 |
| THR | 0.71 | 0.66 | 0.62 | 0.70 |
| PHE | 0.40 | 0.20 | 0.33 | 0.30 |

Table S11: Matrix denoting the effective energy scales (in kcal/mol) for DNA-protein contacts between the sugar group (S) and amino-acid side-chain (SC).

| | GUA | ADE | CYT | THY |
|---|---|---|---|---|
| ALA | 0.56 | 0.44 | 0.47 | 0.45 |
| GLY | 0.80 | 0.79 | 0.73 | 0.73 |
| LEU | 0.06 | 0.07 | 0.01 | 0.11 |
| ILE | 0.31 | 0.33 | 0.25 | 0.24 |
| TYR | 0.62 | 0.58 | 0.61 | 0.57 |
| TRP | 0.38 | 0.45 | 0.45 | 0.41 |
| LYS | 0.97 | 0.90 | 0.89 | 0.89 |
| PRO | 0.49 | 0.50 | 0.49 | 0.50 |
| GLU | 0.32 | 0.26 | 0.31 | 0.21 |
| HIS | 0.83 | 0.86 | 0.75 | 0.78 |
| ASP | 0.54 | 0.39 | 0.56 | 0.18 |
| ARG | 1.14 | 0.99 | 0.94 | 1.02 |
| GLN | 0.68 | 0.69 | 0.67 | 0.55 |
| ASN | 0.81 | 0.75 | 0.76 | 0.79 |
| CYS | 0.56 | 0.28 | 0.28 | 0.42 |
| SER | 0.94 | 0.75 | 0.86 | 0.78 |
| MET | 0.35 | 0.38 | 0.45 | 0.30 |
| VAL | 0.42 | 0.39 | 0.33 | 0.36 |
| THR | 0.82 | 0.78 | 0.76 | 0.85 |
| PHE | 0.47 | 0.36 | 0.41 | 0.37 |

Table S12: Matrix denoting the effective energy scales (in kcal/mol) for DNA-protein contacts between nucleobase (B) and amino-acid backbone (BB).

| | GUA | ADE | CYT | THY |
|---|---|---|---|---|
| ALA | 0.69 | 0.68 | 0.67 | 0.81 |
| GLY | 1.11 | 1.04 | 1.11 | 1.04 |
| LEU | 0.00 | 0.10 | 0.14 | 0.35 |
| ILE | 0.37 | 0.44 | 0.47 | 0.52 |
| TYR | 0.60 | 0.51 | 0.72 | 0.58 |
| TRP | 0.33 | 0.60 | 0.58 | 0.62 |
| LYS | 0.96 | 0.83 | 1.01 | 0.85 |
| PRO | 0.58 | 0.62 | 0.61 | 0.67 |
| GLU | 0.51 | 0.22 | 0.48 | 0.30 |
| HIS | 1.11 | 0.96 | 1.00 | 1.04 |
| ASP | 0.86 | 0.45 | 0.86 | 0.37 |
| ARG | 1.18 | 1.02 | 1.16 | 1.13 |
| GLN | 0.86 | 0.94 | 0.92 | 0.98 |
| ASN | 1.11 | 1.09 | 1.07 | 1.19 |
| CYS | 0.62 | 0.60 | 0.70 | 0.58 |
| SER | 1.16 | 1.02 | 1.18 | 1.08 |
| MET | 0.38 | 0.59 | 0.66 | 0.68 |
| VAL | 0.62 | 0.58 | 0.56 | 0.65 |
| THR | 0.93 | 0.95 | 0.97 | 1.03 |
| PHE | 0.61 | 0.43 | 0.62 | 0.65 |

Table S13: Matrix denoting the effective energy scales (in kcal/mol) for DNA-protein contacts between nucleobase (B) and amino-acid side-chain (SC).

| | GUA | ADE | CYT | THY |
|---|---|---|---|---|
| ALA | 0.75 | 0.71 | 0.70 | 0.78 |
| GLY | 1.07 | 1.02 | 1.04 | 1.01 |
| LEU | 0.00 | 0.03 | 0.01 | 0.24 |
| ILE | 0.32 | 0.38 | 0.43 | 0.40 |
| TYR | 0.86 | 0.74 | 0.90 | 0.85 |
| TRP | 0.21 | 0.20 | 0.46 | 0.39 |
| LYS | 1.13 | 1.09 | 1.22 | 1.12 |
| PRO | 0.63 | 0.64 | 0.58 | 0.69 |
| GLU | 0.69 | 0.42 | 0.66 | 0.43 |
| HIS | 1.21 | 1.13 | 1.18 | 1.17 |
| ASP | 1.00 | 0.59 | 0.98 | 0.53 |
| ARG | 1.52 | 1.38 | 1.49 | 1.41 |
| GLN | 1.09 | 1.06 | 1.03 | 1.08 |
| ASN | 1.22 | 1.20 | 1.21 | 1.30 |
| CYS | 0.56 | 0.50 | 0.68 | 0.44 |
| SER | 1.27 | 1.11 | 1.24 | 1.14 |
| MET | 0.51 | 0.62 | 0.63 | 0.72 |
| VAL | 0.63 | 0.61 | 0.55 | 0.64 |
| THR | 1.09 | 1.07 | 1.08 | 1.15 |
| PHE | 0.64 | 0.50 | 0.54 | 0.62 |

Table S14: PDB IDs of the DNA sequences used for deriving the coarse-grained form factors for the TIS representation.

122D 123D 183D 1D3R 1DPN 1EDR 1G75 1I3T 1N1O 1P54 1VE8 2D25 403D 436D 454D 456D 460D 158D 196D 1BD1 1BNA 1D23 1D49 1D56 1D8G 1D8X 1DOU 1EHV 1EN3 1EN8 1EN9 1ENE 1ENN 1FQ2 1IKK 1JGR 1L4J 1M6G 1NVN 1NVY 1P4Y 1S23 1S2R 1SGS 1SK5 1UB8 1ZF0 1ZF3 1ZF4 1ZF5 1ZF7 1ZFB 1ZFF 1ZFG 232D 251D 307D 355D 3DNB 423D 428D 431D 455D 476D 477D 5DNB 9BNA

Table S15: Coarse-grained form factors derived using IBA for the phosphate group ($f_1(q)$), non-terminal sugar ($f_2(q)$), 3′ sugar ($f_3(q)$), 5′ sugar ($f_4(q)$), adenine ($f_5(q)$), guanine ($f_6(q)$), thymine ($f_7(q)$), and cytosine ($f_8(q)$).

| q ($nm^{-1}$) | $f_1(q)$ | $f_2(q)$ | $f_3(q)$ | $f_4(q)$ | $f_5(q)$ | $f_6(q)$ | $f_7(q)$ | $f_8(q)$ |
|---|---|---|---|---|---|---|---|---|
| 0.00 | 26.13 | 5.62 | 9.51 | 9.51 | 17.97 | 22.29 | 16.93 | 9.56 |
| 0.10 | 26.13 | 5.62 | 9.51 | 9.51 | 17.97 | 22.29 | 16.93 | 9.56 |
| 0.20 | 26.13 | 5.62 | 9.51 | 9.51 | 17.97 | 22.28 | 16.93 | 9.55 |
| 0.30 | 26.12 | 5.61 | 9.51 | 9.51 | 17.96 | 22.27 | 16.92 | 9.55 |
| 0.40 | 26.12 | 5.61 | 9.50 | 9.50 | 17.95 | 22.25 | 16.91 | 9.54 |
| 0.50 | 26.11 | 5.60 | 9.49 | 9.49 | 17.94 | 22.23 | 16.90 | 9.53 |
| 0.60 | 26.10 | 5.60 | 9.48 | 9.48 | 17.92 | 22.21 | 16.88 | 9.52 |
| 0.70 | 26.08 | 5.59 | 9.46 | 9.47 | 17.90 | 22.18 | 16.86 | 9.51 |
| 0.80 | 26.07 | 5.58 | 9.45 | 9.45 | 17.87 | 22.15 | 16.84 | 9.50 |
| 0.90 | 26.05 | 5.57 | 9.43 | 9.44 | 17.85 | 22.11 | 16.81 | 9.48 |
| 1.00 | 26.04 | 5.56 | 9.41 | 9.42 | 17.82 | 22.07 | 16.79 | 9.46 |
| 1.10 | 26.02 | 5.55 | 9.39 | 9.40 | 17.78 | 22.02 | 16.76 | 9.45 |
| 1.20 | 25.99 | 5.54 | 9.37 | 9.37 | 17.75 | 21.97 | 16.72 | 9.42 |
| 1.30 | 25.97 | 5.52 | 9.34 | 9.35 | 17.71 | 21.92 | 16.69 | 9.40 |
| 1.40 | 25.94 | 5.51 | 9.32 | 9.33 | 17.67 | 21.86 | 16.65 | 9.38 |
| 1.50 | 25.92 | 5.49 | 9.29 | 9.30 | 17.62 | 21.79 | 16.61 | 9.35 |
| 1.60 | 25.89 | 5.48 | 9.26 | 9.27 | 17.57 | 21.73 | 16.57 | 9.32 |
| 1.70 | 25.86 | 5.46 | 9.23 | 9.24 | 17.52 | 21.66 | 16.52 | 9.29 |
| 1.80 | 25.82 | 5.44 | 9.19 | 9.21 | 17.47 | 21.58 | 16.47 | 9.26 |
| 1.90 | 25.79 | 5.42 | 9.16 | 9.17 | 17.41 | 21.50 | 16.42 | 9.23 |
| 2.00 | 25.75 | 5.40 | 9.12 | 9.14 | 17.36 | 21.42 | 16.37 | 9.20 |
| 2.10 | 25.71 | 5.38 | 9.08 | 9.10 | 17.29 | 21.34 | 16.32 | 9.16 |
| 2.20 | 25.67 | 5.36 | 9.04 | 9.07 | 17.23 | 21.25 | 16.26 | 9.12 |
| 2.30 | 25.63 | 5.34 | 9.00 | 9.03 | 17.16 | 21.15 | 16.20 | 9.08 |
| 2.40 | 25.59 | 5.32 | 8.96 | 8.99 | 17.09 | 21.06 | 16.14 | 9.04 |
| 2.50 | 25.55 | 5.30 | 8.91 | 8.94 | 17.02 | 20.96 | 16.07 | 9.00 |
| 2.60 | 25.50 | 5.28 | 8.87 | 8.90 | 16.95 | 20.85 | 16.01 | 8.96 |
| 2.70 | 25.45 | 5.25 | 8.82 | 8.86 | 16.87 | 20.74 | 15.94 | 8.92 |
| 2.80 | 25.40 | 5.23 | 8.78 | 8.81 | 16.79 | 20.63 | 15.87 | 8.87 |
| 2.90 | 25.35 | 5.20 | 8.73 | 8.77 | 16.71 | 20.52 | 15.80 | 8.82 |
| 3.00 | 25.30 | 5.18 | 8.68 | 8.72 | 16.63 | 20.40 | 15.73 | 8.78 |
| 3.10 | 25.24 | 5.16 | 8.63 | 8.67 | 16.55 | 20.29 | 15.65 | 8.73 |
| 3.20 | 25.19 | 5.13 | 8.58 | 8.63 | 16.46 | 20.16 | 15.58 | 8.68 |
| 3.30 | 25.13 | 5.11 | 8.52 | 8.58 | 16.37 | 20.04 | 15.50 | 8.63 |
| 3.40 | 25.07 | 5.08 | 8.47 | 8.53 | 16.28 | 19.91 | 15.42 | 8.57 |
| 3.50 | 25.01 | 5.06 | 8.42 | 8.48 | 16.18 | 19.78 | 15.34 | 8.52 |
| 3.60 | 24.95 | 5.03 | 8.36 | 8.43 | 16.09 | 19.65 | 15.26 | 8.47 |
| 3.70 | 24.89 | 5.01 | 8.31 | 8.38 | 15.99 | 19.51 | 15.17 | 8.41 |
| 3.80 | 24.83 | 4.98 | 8.25 | 8.33 | 15.90 | 19.37 | 15.09 | 8.36 |
| 3.90 | 24.76 | 4.96 | 8.20 | 8.28 | 15.80 | 19.23 | 15.01 | 8.30 |
| 4.00 | 24.70 | 4.93 | 8.14 | 8.22 | 15.69 | 19.09 | 14.92 | 8.24 |
| 4.10 | 24.63 | 4.91 | 8.08 | 8.17 | 15.59 | 18.95 | 14.83 | 8.18 |
| 4.20 | 24.56 | 4.89 | 8.03 | 8.12 | 15.49 | 18.80 | 14.75 | 8.12 |
| 4.30 | 24.49 | 4.86 | 7.97 | 8.07 | 15.38 | 18.65 | 14.66 | 8.06 |
| 4.40 | 24.42 | 4.84 | 7.91 | 8.02 | 15.27 | 18.50 | 14.57 | 8.00 |
| 4.50 | 24.35 | 4.82 | 7.86 | 7.96 | 15.17 | 18.35 | 14.48 | 7.94 |
| 4.60 | 24.27 | 4.80 | 7.80 | 7.91 | 15.06 | 18.20 | 14.39 | 7.88 |
| 4.70 | 24.20 | 4.78 | 7.74 | 7.86 | 14.95 | 18.04 | 14.30 | 7.82 |
| 4.80 | 24.12 | 4.76 | 7.69 | 7.81 | 14.83 | 17.88 | 14.20 | 7.76 |
| 4.90 | 24.04 | 4.74 | 7.63 | 7.76 | 14.72 | 17.73 | 14.11 | 7.69 |
| 5.00 | 23.97 | 4.72 | 7.57 | 7.71 | 14.61 | 17.57 | 14.02 | 7.63 |
